# *In situ* quantification of ribosome number by electron tomography

**DOI:** 10.1101/2024.09.27.614074

**Authors:** Mounir El Hankouri, Marco Nousch, Thomas Müller-Reichert, Gunar Fabig

## Abstract

Ribosomes, discovered in 1955 by George Palade, were originally described as small cytoplasmic particles preferentially associated with the endoplasmic reticulum (ER). Over the years, numerous studies were conducted on both ribosome structure and function. The basic question, however, how many ribosomes are contained within whole cells has not been addressed so far. Here we developed a microscopic method to determine the total number of ribosomes in human hTRET-RPE-1 and in nematode cells from different tissues in *Caenorhabditis elegans* hermaphrodites. Electron tomography of high-pressure frozen, freeze substituted and resin-embedded samples revealed that the number of ribosomes as determined by this electron tomographic approach closely matches the biochemical estimates for the hTRET-RPE-1 cells. In addition, RPOA-1-depleted worms compared to control worms showed an expected decrease in the number of ribosomes in two out of three different tissue types investigated. The application of our imaging-based approach to directly count the number of ribosomes in given samples will significantly enhance our basic understanding of ribosome localization and distribution within cells and tissues.

## Introduction

Ribosomes are the cellular sites of protein synthesis. These macromolecular machines translate messenger RNA (mRNA) into polypeptide chains, thus facilitating the expression of genes into functional proteins.^1^ Emphasizing the key role of ribosomes in regulating cellular metabolism, research in the past was mainly focused on their translational activity. All ribosomes are composed of two subunits, a large and a small subunit. The large 60S subunit of the eukaryotic ribosome consists of three ribosomal RNA (rRNA) molecules (28S, 5.8S, and 5S in mammals) and 46 proteins.^2^ The small 40S subunit consists of one rRNA chain (18S) and 33 proteins. Of the total 79 proteins in a ribosome, 32 have no structural homologs to either bacterial or archaeal ribosomes, and those that do have homologs still harbor large eukaryote-specific extensions.^3^ Apart from the variability in certain rRNA expansion segments, all eukaryotic ribosomes, from yeast to human cells, are very similar in structure.^4^ Despite our knowledge on ribosome function and structure in both prokaryotes and eukaryotes, the analysis of the total number of ribosomes in cells has not been studied in detail.

In practice, ribosome number can be analyzed by either measuring the rRNA content (e.g., by capillary electrophoresis),^5^ or by quantifying either ribosomal protein content via Western blot or fluorescence intensity.^6,7^ Both methods rely on indirect measurements. An extrapolation of the ribosome number often gives inaccurate estimates, obvious when comparing results obtained by both methods.^8^ To overcome these limitations, we explored electron tomography as an imaging-based method for ribosome quantification. We established a semi-automatic image analysis pipeline to segment and count ribosomes in mammalian hTERT-RPE-1 cells. In addition, we also analyzed ribosome number in the gonadal region and in somatic tissue (vulva) of the nematode *Caenorhabditis elegans*. For *C. elegans*, we further aimed to compare control and RNAi-treated worms with an expected difference in the number of ribosomes. For this, we chose to knock-down the RPOA-1 (RNA polymerase I A subunit) via RNAi feeding. Here, we show that transmission electron microscopy, particularly electron tomography, together with our newly established image analysis pipeline for semi-automatic segmentation of particles can be used to determine the number of ribosomes in both single cells and in cells organized in complex tissues. Our 3D method is suitable for a robust quantification of ribosome number and can be applied to a broad range of model systems as ribosomes can be reliably detected in almost all samples using conventional transmission electron microscopy techniques.9–14

## Materials and methods

### Culture of hTERT-RPE-1 cells

The hTERT-RPE-1cells (ATCC CRL-4000, Manassas, USA) were grown in 750 ml-cell culture flasks (Greiner Bio-One GmbH, Frickenhausen, Germany) containing 10 ml of DMEM medium (DMEM Glutamax, Gibco Fischer Scientific, Schwerte, Germany) with 10% (w/v) FBS (Merck KGaA, Darmstadt, Germany) and 1x (v/v) penicillin/ streptomycin antibiotics (Gibco Fischer Scientific, Schwerte, Germany). Cells were kept in a humidified incubator supplemented with 5% CO_2_ at 37°C and passaged every 2-3 days to avoid full confluency of the culture. For this, the old medium was discarded, and the cells were washed with pre-warmed phosphate buffered saline (PBS, Gibco Fischer Scientific, Schwerte, Germany). The hTERT-RPE-1 cells were then treated with 0.25% trypsin-EDTA (Gibco Fischer Scientific, Schwerte, Germany) for 3 min to detach the cells. Subsequently, about 10 ml volume of cells was transferred to a new cell culture flask and re-suspended in fresh DMEM medium.

### High-pressure freezing and freeze substitution

The hTERT-RPE-1 cells were attached to Sapphire discs as previously published.^15^ Briefly, a ‘sandwich’ for high-pressure freezing was assembled by gently placing a Sapphire disc (Wohlwend, article no. 500) upside down on a pre-wetted aluminum planchette with a cavity of 0.04 mm (Wohlwend, article no. 737). As a cryo-protectant we used the cell culture medium supplemented with additional 10% (w/v) BSA (Carl Roth, Karlsruhe, Germany). Samples were cryo-immobilized at a pressure of ∼2000 bar and a cooling rate of ∼20,000°C/sec using a Compact 03 (Wohlwend, Switzerland) high-pressure freezer as published.^16,17^ For cryo-immobilization of worms, sample holders (type-A aluminum planchettes; Wohlwend, article no. 241) were pre-wetted with hexadecene and then filled with 10% (w/v) polyvinylpyrrolidone (PVP) (MW 10,000, Merck KGaA, Darmstadt, Germany) diluted in M9 buffer. About five gravid adult hermaphrodites (24 hours after L4 stage) were transferred to the 100 µm-indentation of a type-A carriers for each round of freezing. The specimen holders were gently closed by placing the flat side of a type-B aluminum planchette (Wohlwend, article no. 242) on top of the type-A planchette. After high-pressure freezing, such closed sample holders were stored in liquid nitrogen until further use.

For the following freeze substitution, sample holders containing either hTERT-RPE-1 cells or worms were opened under liquid nitrogen with a syringe needle. Sapphire discs with attached frozen hTERT-RPE-1 cells and type-A planchettes containing worms were transferred to cryo-vials filled with anhydrous acetone containing 1% (w/v) osmium tetroxide (EMS) and 0.1% (w/v) uranyl acetate (Polysciences, Warrington, UK) as previously published.^18^ Freeze substitution was carried out using an automatic freeze substitution machine (AFS2, Leica Microsystems, Vienna, Austria). Samples were kept at -90°C for one hour, warmed up to -30°C with increments of 5°C/hr, kept for 5 hrs at -30°C and then warmed up to 0°C in steps of 5°C/h.^17^

### Resin embedding, re-mounting and ultramicrotomy

Directly after the freeze-substitution, hTERT-RPE-1 and worm samples were washed three times with pure anhydrous acetone and infiltrated with Epon/Araldite (EMS, Hatfield, USA) at increasing concentrations of resin (resin:acetone: 1:3, 1:1, 3:1, then pure resin) for 2 hrs each step at room temperature.^18^ Samples were incubated with pure resin overnight and then again for 4 hrs. hTERT-RPE-1 cells were mounted in flow-through chambers as published.^16^ Worms were thin-layer embedded between two Teflon-coated glass slides as described.^18^ Samples were polymerized at 60°C for 48 hrs. hTERT-RPE-1 cells in resin blocks and the thin-layer embedded worms re-mounted on dummy blocks^18^ were serially sectioned using an ultramicrotome (EM UC6, Leica Microsystems, Austria). Ribbons of serial (300 nm) sections were collected on Formvar-coated copper slot grids and post-stained with 2% (w/v) uranyl acetate in 70% (v/v) methanol followed by 0.4% (w/v) lead citrate in double-distilled water.^16^ Finally, both sides of the samples were coated with colloidal gold (20 nm; BBI, UK) serving as fiducial markers for subsequent tomographic reconstruction.^15^

### Electron tomography and 3D reconstruction

Serial semi-thick sections (300 nm) were pre-screened at low magnification by using a transmission electron microscope (Morgagni, Thermo Fisher Scientific) operated at 80 kV and equipped with a 2k x 2k CCD camera (Veleta, EMSIS). This allowed to locate hTERT-RPE-1 cells in interphase. We also identified distal and pachytene regions and also vulva cells in *C. elegans* samples. Sections were then transferred to a transmission electron microscope (Tecnai F30, Thermo Fisher Scientific, USA) operated at 300 kV and equipped with a 4k x 4k CMOS camera (OneView, Gatan, USA). Using a dual-axis specimen holder (Type 2040, Fischione Instruments, USA), tilt series were recorded from -60° to +60° with 1° increments at a magnification of 4,700x and a pixel size of 2.572 nm using the SerialEM software package.^19,20^ Subsequently, grids were rotated for 90° in the X/Y-plane and second tilt series were acquired using identical microscope settings.^21^ For each region of interest, the tomographic A- and B-stack of was then reconstructed, combined and flattened using the IMOD software package.^22,23^

### Automatic segmentation of ribosomes in electron tomograms

Six tomograms each of six different hTERT-RPE-1 cells and four tomograms each of the distal gonadal region, the pachytene region and the vulva in four control animals and three tomograms each of the distal gonadal region, the pachytene region and the vulva in three *rpoa-1(RNAi)* animals were generated. The MRC-files were then converted to TIFF-files and further processed with the Fiji software package (version 1.8.0_322).^24^ This program allowed a selection of regions of interest (ROIs) and was used for cropping out selected sub-regions of interest from each tomogram devoid of any organelles such as mitochondria. All selected regions and the accompanying tomograms are listed in Table 1 and can be accessed via a publicly available [OMERO link](https://omero.med.tu-dresden.de/webclient/?show=project-1303).

These TIFF-files were then imported in Arivis Vision4D 3.4. After setting the pixel size to 2.57 nm, the following procedure was applied: as an input, all planes of the cropped image were selected (ROI: custom → planes → all → scaling 100%). The images were inverted to detect the stained ribosomes as local maxima and then a background correction algorithm was applied (background correction → preserve bright → blur diameter = 386 nm) to remove background noise. The resulting images were stored. Next, the segmentation function “blob finder” was used to segment the ribosomes. The specific parameters for the analyzed images were: diameter, 20 nm; probability threshold, 30% and split sensitivity, 70%. Because specimens shrink due to dehydration during preparation and a sample collapse in the electron beam during the acquisition of tomograms,^25,26^ we chose a value of 20 nm for the diameter of the ribosomes. This value approximates the lower bound of the average diameter of eukaryotic ribosomes, which is reported to be between 25 and 30 nm.^27,28^ In addition, the probability threshold was optimized empirically to automatically segment ribosomes showing a relatively low contrast.

After application of the “blob finder” algorithm, a list of possible segmented ribosomes was created. The automatically segmented ribosomes in the first and last slices of the tomogram were deleted via the objects table because these tomographic slices have usually a very bad contrast which could lead to erroneous detected objects. In addition, all segmented objects in only a single plane and all with a voxel count lower three were removed. The colloidal gold particles, used as fiducials during tomographic reconstruction at the section boundaries, were also manually removed. All objects with a voxel number of three or above were checked if they matched a ribosome. For that, the tomograms of the selected regions were opened in Fiji. The identical plane with the putative segmented ribosomes was displayed next to it in Arivis, and by comparing the intensity and shape in the previous and following slices, the segmented objects were checked.

### Calculation of the total number of ribosomes in hTERT-RPE-1 cells

In previous publications we calculated a shrinkage factor to account for the sample collapse that occurred during acquisition of the tomographic data.^11^ This factor was given by the ratio between the calculated thickness (as obtained after tomographic reconstruction of stitched serial sections) and the initial thickness (as obtained from the ultramicrotome settings and number of sections). This factor was then multiplied with the pixel size in the Z-dimension of each tomogram to account for the shrinkage. However, in this work we rely on single, semi-thick sections through given samples and hence we were much more prone to fluctuations of physical section thickness from ultramicrotomy. As we still wanted to account for the shrinkage of this single tomographic section, we collected a comprehensive list of 77 stitched data sets, both published and unpublished, that have been created in our lab (see Suppl. Data 5) and determined a mean shrinkage factor of 1.575. This factor we applied to all our current data sets to correct for sample shrinkage in the Z-dimension. After cropping five sub-regions per tomogram to segment the ribosomes in them, we calculated their volume by multiplying the X, Y and Z dimensions. The number of objects per sub-region volume was then calculated per cubic micrometer and divided by the shrinkage factor. This value represents the normalized, shrinkage-corrected total number of ribosomes per cubic micrometer of cytoplasmic volume. An identical calculation was performed to five cropped sub-regions per tomogram in distal and pachytene regions of the gonad and also in somatic cells of the vulva in control worms. A similar analysis was performed in *rpoa-1(RNAi)* worms.

### Determination of hTERT-RPE-1 cell size

The volume of hTERT-RPE-1 cells was determined using cells (up to passage no. 4) treated with 0.25% trypsin followed by a gentle pipetting to break cell-cell contacts. Subsequently, cells were incubated in a cell imaging dish for 2 - 4 hours to allow attachment, staining, and fixation, ensuring round cells devoid of cell-cell contacts. Staining involved MemBrite® Fix Pre-Staining Solution 1X (Biotium, Fremont, USA) followed by MemBrite® Fix 543/560 dye solution 1X (Biotium, Fremont, USA), with subsequent fixation in 100% ice-cold methanol. The nuclei were stained with DAPI (Merck KGaA, Darmstadt, Germany). Such fixed and stained cells were analyzed by using a Leica TCS SP5 laser scanning confocal microscope (Leica Microsystems, Wetzlar, Germany) with a 40x/1.25 HCX PL APO lambda blue oil objective. Initially, cells were located through the ocular before transitioning to laser scanning for image acquisition. For the DAPI staining, samples were illuminated with a 405 nm laser and signals were collected within the spectral ranges of 415 - 480 nm. For the MemBrite staining, samples were illuminated with a 543 nm laser and signals were collected within the spectral ranges of 550 - 700 nm. Image were acquired in a XY-scan field of 512 x 512 pixels and with a Z-dimension big enough to capture the entire cell volume (40-50 µm).

The images obtained by laser scanning confocal microscopy underwent initial processing using Fiji, which consisted first in merging the fluorescence channels (DAPI and MemBrite dye) of each image, then pre-selecting the cells of interest and cropping them for individual analysis. The cropped images were then saved as TIFF-files, imported in Arivis Vision4D 3.5 and further analyzed. In the Arivis analysis pipeline the bilateral denoising (diameter 5 µm, sensitivity 51) and the automatic background correction (blur diameter 150 µm, preserve bright) algorithms were used. The entire image stack of the MemBrite-dye channel was chosen for membrane segmentation using the magic wand tool, enabling precise segmentation of cell boundaries across all planes. Objects were then manually merged into 3D objects in the object table to facilitate analysis of the cell volume. Subsequently, the same methodology was applied to segment nuclei in the DAPI channel and determine their volumes. For each of ten cells in interphase, the volume of nucleus, where no ribosomes are present, was subtracted from the volume of the entire cell. The determined average of the cell volume was then taken for further calculations.

### High-resolution automated electrophoresis of RNA samples

The hTERT-RPE-1 cells at a confluency level of about 80% were counted by transferring 2 x 5 μl of cell suspension into a hemocytometer. These initial cell cultures were diluted to obtain triplicates of suspensions containing each either one or two million cells in 15 ml Falcon tubes. The tubes were centrifuged for 6 min at 16,000 rcf to sediment the cells. The medium was then removed and the tubes containing the pellets were immediately placed in liquid nitrogen to fix the cells. The total RNA was then extracted using a previously published phenol:chloroform protocol.^29^ The total RNA was eluted in 50 µl of RNase-free water and then diluted 1:50 in RNase-free water to reach a concentration which could be reliably measured.

### Extrapolation of number of ribosomes from RNA capillary electrophoresis data

The quality and quantity of the total RNA, including the 18S and 28S ribosomal RNA fragments extracted from the two cell concentrations of hTERT-RPE-1 (one and two million cells), were assessed using an Agilent 2100 Bioanalyzer (Agilent, Santa Clara, USA) and Agilent RNA 6000 Nano Kit (Agilent, Santa Clara, USA) according to the manufacturer’s protocol (https://www.agilent.com/cs/library/usermanuals/Public/G2938-90034_RNA6000Nano_KG.pdf). In a successful run, the software displays a data plot of size/migration time *versus* fluorescence intensity. Peaks were automatically identified and tabulated by peak ID, including ribosomal RNA fragments (see Suppl. Data 6).

An RNA ladder containing a mixture of RNA of known concentrations was run first. To calculate the total RNA concentration, the area under the entire RNA electropherogram was automatically determined. The ladder, which provided the concentration-to-area ratio, was used by the software to transform area values into concentration values. For the 18S and 28S rRNA fragments, the areas under their respective peaks were automatically determined. A threshold or baseline under the peaks was set by the software to measure the area. Due to potential inaccuracies in the automated thresholds or baselines, a manual adjustment was applied as recommended by the manufacturer’s user guide.^30^

The area under the 18S and 28S rRNA peaks was then converted to concentrations. The RNA integrity number (RIN) was automatically determined and displayed in the results sub-tab and below the gel-like image. The RIN generated by the bioanalyzer indicated rRNA quality, with values between 7 and 10 denoting good quality.^31^ Only samples with RIN > 9 were considered for further calculations.

To determine the number of ribosomes per hTERT-RPE-1 cell, we calculated the number of the 18S and 28S rRNA subunits using the equation provided below (see also Suppl. Data 3). We then determined the mean of the total numbers of these two subunits. Since one ribosome consists of one 18S and one 28S subunit, the mean number of these subunits per cell represents the final number of ribosomes per hTERT-RPE-1 cell.

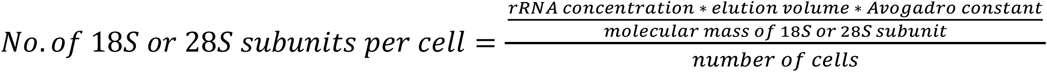

### Cultivation of C. elegans

*C. elegans* (Bristol, N2) hermaphrodites were cultivated on nematode growth medium (NGM) agar plates spotted with *E. coli* OP50 according to established protocols.^32^ For worm propagation, ten worms were transferred to fresh NGM plates every three days.

### RNAi of RPOA-1 in C. elegans

RNA-mediated interference (RNAi) was carried out by feeding bacteria expressing double-stranded RNA according to published protocols.^33,34^ The RNAi feeding construct targeting the RPOA-1 transcript was generated by cloning nucleotides 4073-5115 from the transcript Y48E1A.1a.1 into the vector pL4440 (Addgene, Plasmid #1654), which was subsequently transformed into *E. coli* HT115(DE3). Expression of dsRNA was induced by the addition of IPTG (Thermo Fisher Scientific GmbH, Dreieich, Germany) to a final concentration of 1 mM as previously described.^35^ About 40 animals in the L4 larval stage were transferred to RNAi-expressing bacteria on NGM plates for 24 h at 25°C. The sequence of the primers used for amplifying nucleotides 4073-5115 from Y48E1A.1a.1 were: primer forward: TGCACTTGTTCGTCGATCCA; primer reverse: GAGAAGACGGTCCTGCAACA.

The genome of wild-type *C. elegans* (N2, Bristol) was used for the amplification of the nucleotides 4073-5115 from the transcript Y48E1A.1a.1.

As a control condition we used the empty vector pL4440, transformed in the similar bacterial strain *E. coli* HT115(DE3) and treated similarly with IPTG as the dsRNA-expressing bacteria.

## Results

### Concentration of ribosomes in sub-volumes of hTERT-RPE-1 cells

We aimed to show in a prove-of-concept study that an image-based approach can lead to a robust quantification of ribosome number in human tissue culture cells. For this, the number of ribosomes in defined cytoplasmic sub-volumes of high-pressure frozen, freeze substituted and resin-embedded hTERT-RPE-1 cells was determined by electron tomography. In total, we generated six tomograms from different hTERT-RPE-1 cells in interphase and selected five sub-volumes from each tomogram for further analysis (Fig. 1A-D). On purpose, we selected tomographic sub-volumes which contained only ribosomes but no other organelles, such as mitochondria or large vesicles. Correcting for background noise and excluding particles smaller than 20 nm (as described in the methods section), we then segmented ribosomes in these sub-volumes using the Arivis Vision4D software package (Fig. 1E-G). Visual inspection confirmed the accuracy of this automatic segmentation procedure. We then calculated the number of ribosomes in the given tomographic volumes. We noticed some variation in the determined number of ribosomes. To correct for differences in the thickness of the acquired tomograms, which might have been induced during ultramicrotomy and/or subsequent exposure of the sections to the electron beam (Fig. 1H), we multiplied the Z-voxel size with a mean shrinkage factor to compensate for the beam-induced shrinkage (see methods section for details). After this correction step had been applied, we detected on average 18,605 ± 4,129 ribosomes per cubic micrometer in these tomograms (Fig. 1I, horizontal line, Suppl. Data 1). Thus, our approach enabled a precise quantification of the number of ribosomes within a given small cytoplasmic sub-volume of hTERT-RPE-1 cells.

**Fig. 1.**
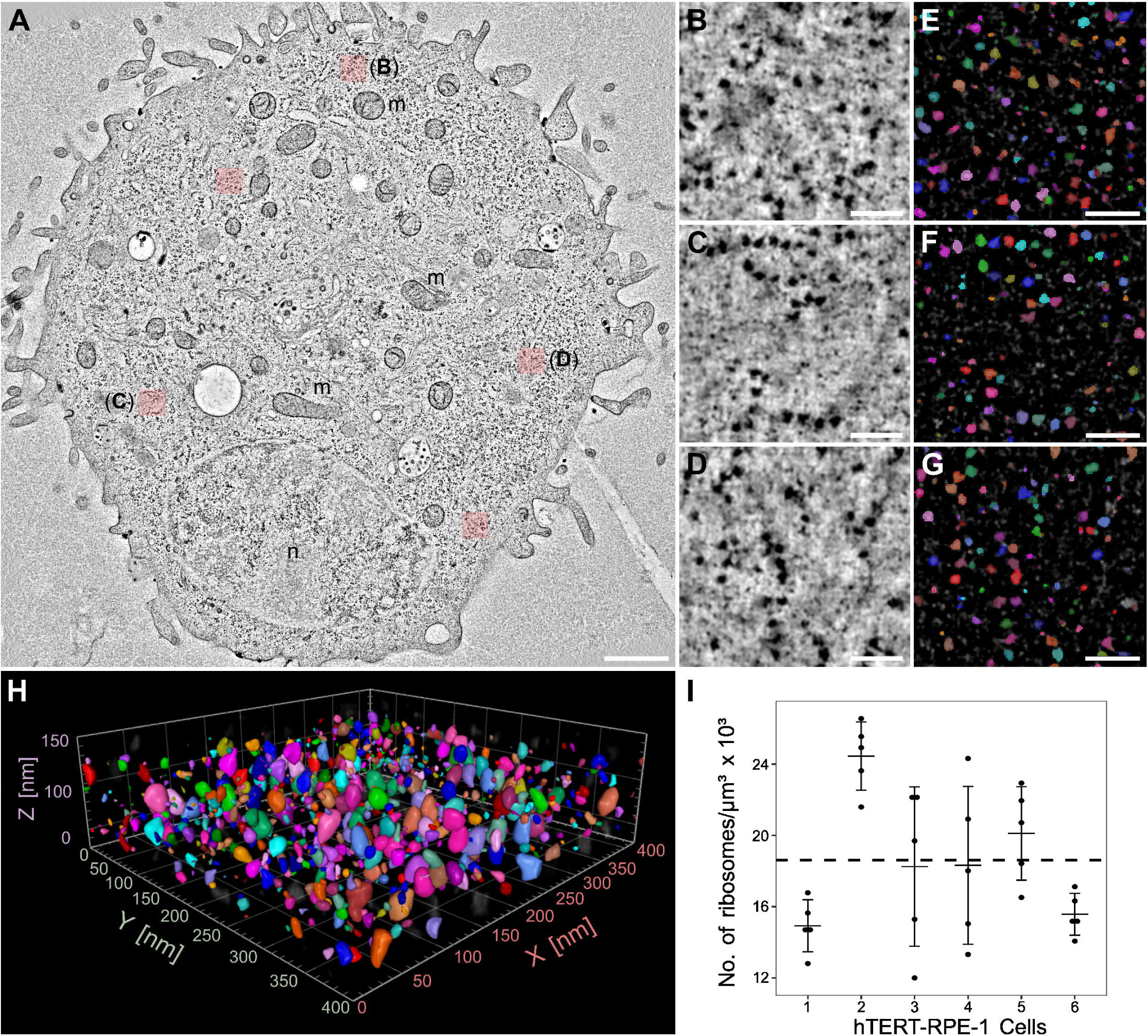
Quantitative analysis of ribosome number in hTERT-RPE-1 cells. (**A**) Tomographic slice showing an [interphase cell](https://omero.med.tu-dresden.de/webclient/?show=dataset-1717) with five selected regions cropped out for detailed data analysis (red overlay). The nucleus (n) of this cell and numerous mitochondria (m) are visible. Scale bar, 1 µm. (**B-D**) Three out of five segmented and analyzed cytoplasmic regions as indicated in (A). Scale bars, 100 nm. (**E-G**) Automatic segmentation of ribosomes in the three selected regions as shown in (B-D). Scale bars, 100 nm. (**H**) Volumetric view showing the ribosomes in the fully segmented Z-stack of the region as shown in (C). (**I**) Quantification of ribosome number in five selected sub-regions of six different hTERT-RPE-1 cells. The horizontal dashed line indicates the mean over all six samples. For the individual plots, the black dots represent the individual measurements from the five sub-regions per cell, the horizontal line indicates the mean and the whiskers show the standard deviation.

### Total number of ribosomes in hTERT-RPE-1 cells via combination of ET and light microscopy approach

Next, we set out to determine the total number of ribosomes in entire hTERT-RPE-1 cells. For this, it was necessary to determine the total cytoplasmic volume of the previously mentioned cells. This was achieved by measuring the volume of individual hTERT-RPE-1 cells by confocal fluorescence microscopy and then subtracting the volume of the nucleus from the total cellular volume. We used MemBrite® to stain the cell membrane and DAPI for staining of the nucleus (Fig. 2A-C). The volumes of ten hTERT-RPE-1 cells and their nuclei were then determined by 3D segmentation using Arivis Vision4D 3.5 (Fig. 2D-F). We observed that the hTERT-RPE-1 cells on average had a total volume of 2,638.9 ± 925.9 μm³ and a nuclear volume of 652.9 ± 137.9 μm³ in interphase. As a result, the hTERT-RPE-1 cells had an average calculated cytoplasmic volume of 1,986 ± 855.4 μm³ (see Suppl. Data 2). We then multiplied this calculated value by the number of ribosomes per 1 μm³ as determined by electron tomography, resulting in an estimated number of 36.9*10^6^ ± 3.5*10^6^ ribosomes per hTERT-RPE-1 cell. Thus, by combining electron and light microscopic analyses, we were able to estimate the total number of ribosomes in the human cell line hTERT-RPE-1.

**Fig. 2.**
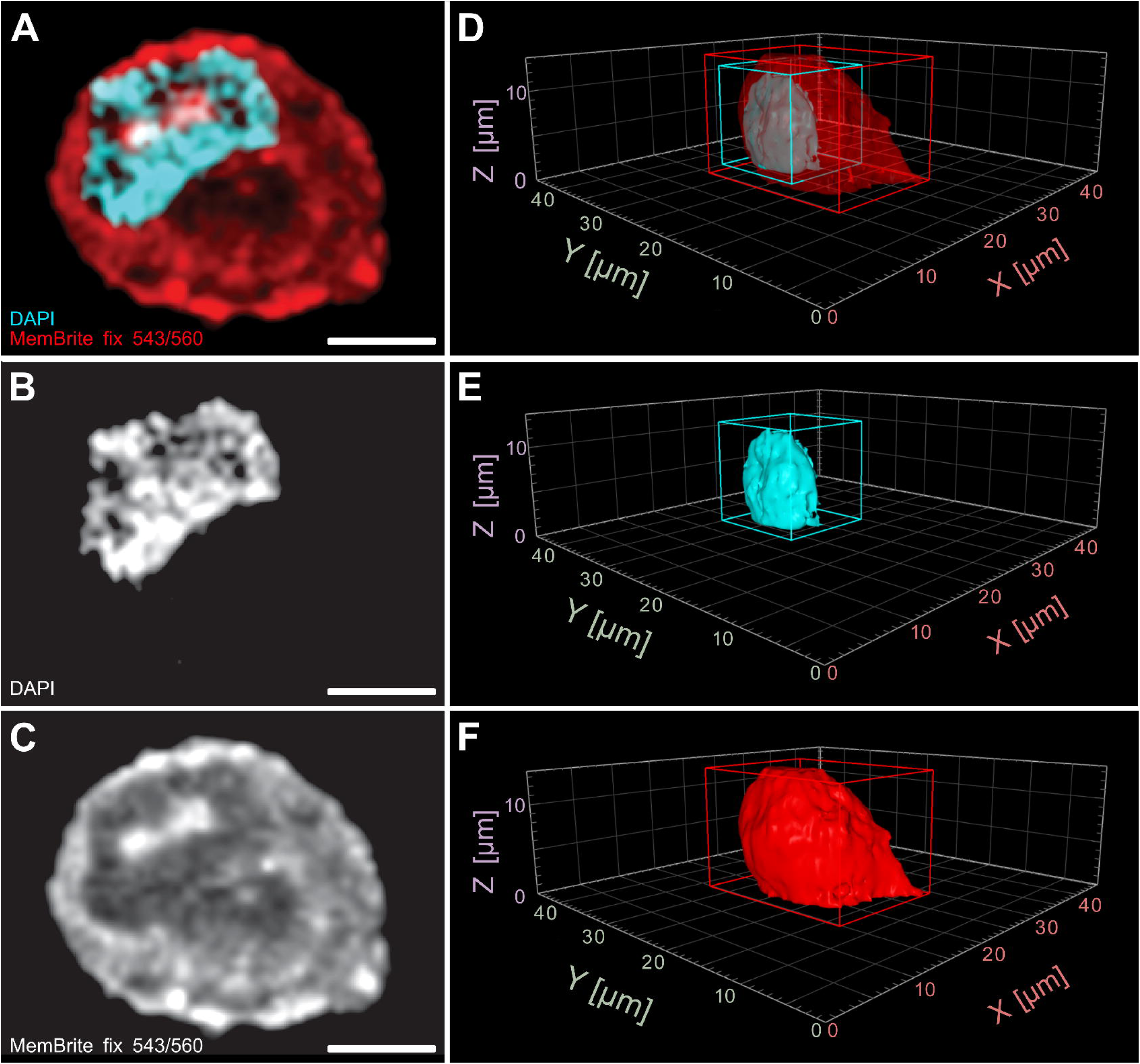
Volume analysis of hTERT-RPE-1 cells by confocal light microscopy. (**A**) Single slice from an acquired confocal Z-stack of a fixed hTERT-RPE-1 cell in interphase. The cell membrane and the cytoplasm are shown in red, the nucleus in cyan. (**B**) Single slice showing only the nucleus in the confocal Z-stack as illustrated in (A). (**C**) Single slice showing only the membrane and cytoplasm in the confocal Z-stack as shown in (A). (**D-F**) Volumetric view of the 3D-segmented cell, nucleus and cytoplasm as shown in (A-C). Scale bars, 5 µm.

### Biochemical quantification of ribosomes in hTERT-RPE-1 cells

We also quantified ribosome number in hTRET-RPE-1 cells using a biochemical approach, thus allowing a direct comparison of biochemical and EM-derived results on ribosome number. We applied capillary electrophoresis on extracted total RNAs from one million and two million hTRET-RPE-1 cells, each in triplicates (Fig. 3A), to ensure accuracy and reproducibility. Using two different cell concentrations validated the results across different scales, confirming that the quantification method is robust and not concentration-dependent. Triplicates provided statistical reliability, reducing experimental variability and ensuring consistent findings.

**Fig. 3.**
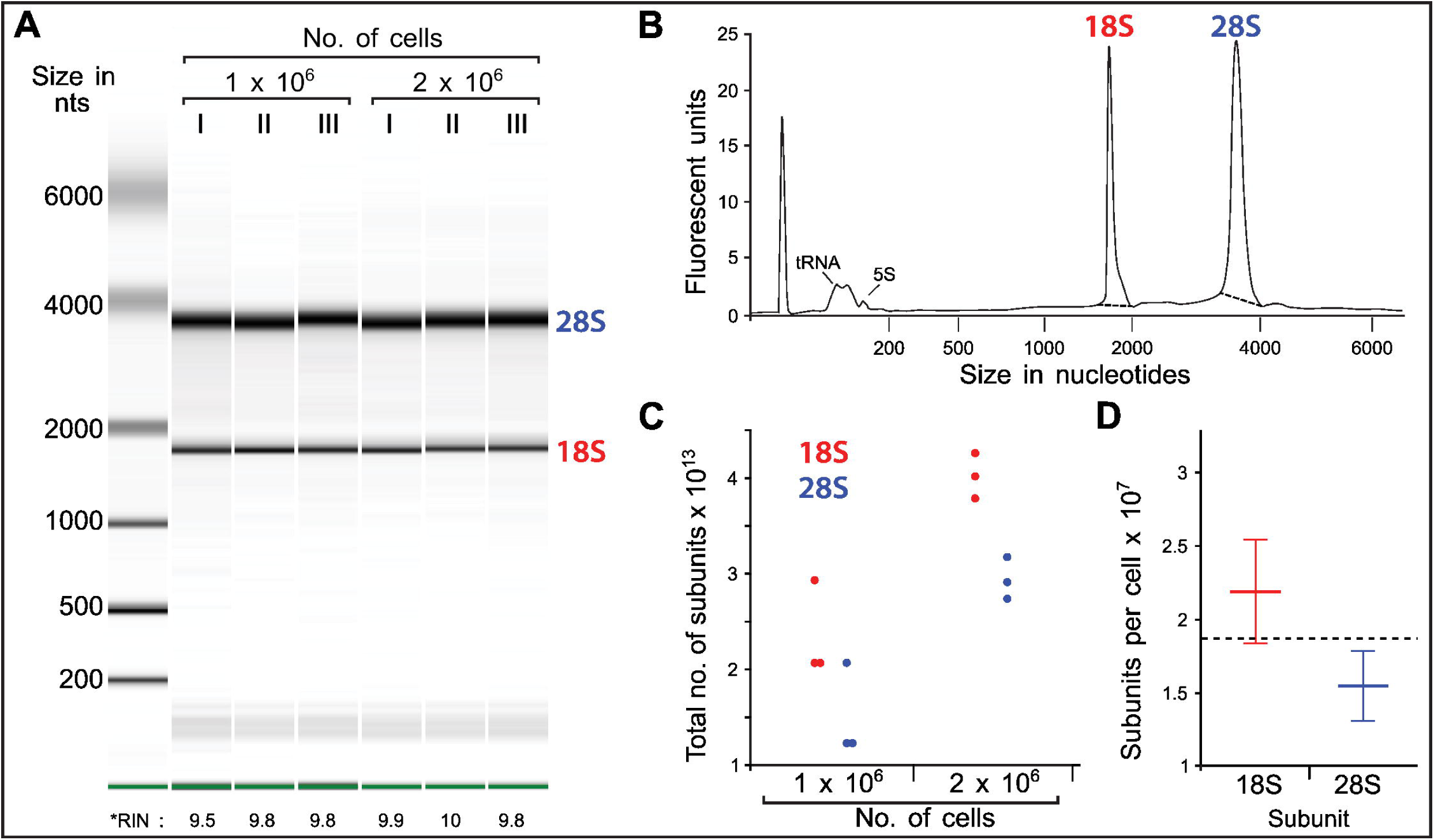
Biochemical determination of ribosome number in hTERT-RPE-1 cells. (**A**) Capillary electrophoresis of extracted total RNA. Green bands at the bottom represent the marker, the bands below 200 nucleotides represent tRNA and 5S RNA, the bands below 2000 nucleotides represent the 18S rRNA, the bands around 4000 nucleotides represent the 28S rRNA. RNA integrity number (RIN) indicates the RNA quality and is given below the plot. (**B**) Example electropherogram of the capillary electrophoresis of the total RNA extracted from one sample (lane II in (A)). (**C**) Graph showing the extrapolated results from the capillary electrophoresis approach estimating the number of subunits of 18S (red) and 28S (blue) subunits in one million (left) and two million (right) cells. (**D**) Graph showing the mean number of 18S and 28S subunits per single hTERT-RPE-1 cell. Error bars indicate the standard deviation. See also Suppl. Data 6.

Only RNA samples with a RNA Integrity Number (RIN) > 9 were considered (Fig. 3A), showing a clean electropherogram with clear 18S and 28S rRNA fragment peaks and no fluctuations in the fluorescence signal (Fig. 3B). Automated measurements of the areas under the 18S and 28S rRNA fragment peaks, using manually adjusted thresholds (indicated by dashed lines in Fig. 3B and colored lines in Suppl. Data 6) were used to calculate the concentration of both rRNA subunits for each sample shown in the digital electrophoresis gel. These concentrations were then converted to the number of subunits (Fig. 3C) using the equation described in the methods section (see also Suppl. Data 3).

The total number of 18S and 28S subunits extracted from two million hTRET-RPE-1 cells was approximately twice that extracted from one million cells (Fig. 3C), validating the robustness of this biochemical quantification approach. To determine the number of ribosomes per hTRET-RPE-1 cell, the number of 18S and 28S subunits was extrapolated. The number of subunits per cell was calculated as the average of the number of 18S and 28S subunits per one million cells and per two million cells, each divided by the respective number of cells (Fig. 3D). In conclusion, given that the total number of 18S and 28S subunits should equal the total number of ribosomes (since the two subunits form one ribosome, as described by Ben-Shem, et al., 2011^4^) the biochemical approach showed an average of 18.7*10^6^ ± 4.4*10^6^ ribosomes per hTRET-RPE-1 cell (Fig. 3D, Suppl. Data 3).

### Quantification of ribosome numbers in C. elegans samples

We further aimed to determine the number of ribosomes in different cell types of the nematode *Caenorhabditis elegans* as this nematode has a relatively small size and a simple body plan (Fig. 4A), allowing us to investigate multiple tissue types in a single semi-thick section for transmission electron microscopy. Furthermore, this animal is well known to be manipulated easily by RNA-mediated interference.^34^ We decided to analyze ribosome number in both, germ cells and somatic cells in four control hermaphrodites prepared for electron microscopy. In more detail, we acquired one tomogram each from cells in the distal (Fig. 4B) and the pachytene (middle) region of the gonad (Fig. 4C), and in one somatic cell of the vulva (Fig. 4D). Specifically, we determined an average of 30,864 ± 4,624 ribosomes in a volume of 1 μm³ in distal cells of the gonad, 27,056 ± 9,197 ribosomes in cells in pachytene. In addition, we detected an average of 18,124 ± 6,797 ribosomes per μm³ in the somatic cells of the vulva (see also Suppl. Data 4). Thus, our analysis revealed that the number of ribosomes per µm³ is not uniform in cells of different tissue origin but varies in different cell types. Interestingly, the density of ribosomes in the somatic cell in the worm is pretty close to the somatic human cell analyzed earlier (Fig. 1I). In conclusion, electron tomography is suitable to detect differences in ribosome number in various tissues of whole organisms.

**Fig. 4.**
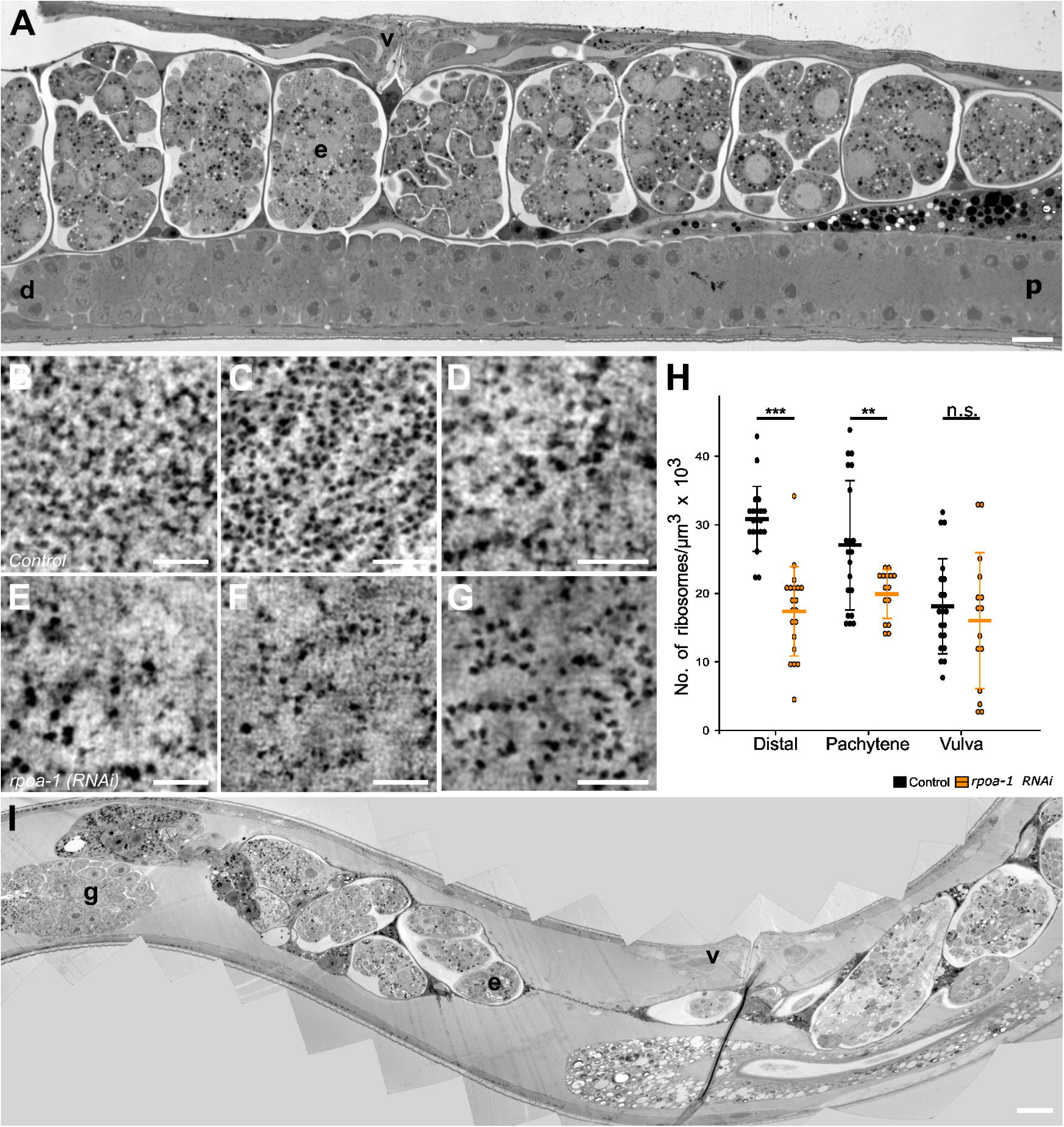
Ribosome number in germ cells and somatic cells of *C. elegans*. (**A**) Overview TEM image showing the morphology of a wild-type hermaphrodite with mitotic germ cells in the distal region of the gonad (d), meiotic cells in the pachytene stage (p, middle region of the gonad), early embryos (e) and somatic cells of the vulva (v). Scale bar, 10 µm. (**B**) Example of tomographic slice from control RNAi *C. elegans* from the distal region of the gonad. (**C**) Pachytene region of the gonad of a control animal. (**D**) Somatic vulva cell of a control animal. Scale bars, 100 nm. (**E**) Examples of tomographic slices from *rpoa-1(RNAi) C. elegans* from the distal region of the gonad. (**F**) Pachytene region of the gonad of a *rpoa-1(RNAi)* animal. (**G**) Somatic vulva cell of a *rpoa-1(RNAi)* animal. Scale bars, 100 nm. (**H**) Number of ribosomes per cubic micrometer of cytoplasm in the distal region, the pachytene region and somatic (vulva) cells in control and *rpoa-1(RNAi)* worms. The single measurements are shown as dots, the mean is represented by a horizontal line and the whiskers indicate the standard deviation. After performing Welch’s two-sided t-test, the resulting levels of significance in between control and test condition are indicated (***: p < 0.001; **: p < 0.01; n.s.: not significant). (**I**) Overview TEM image of a *rpoa-1(RNAi)* worm, showing the vulva (v), the gonad (g) and embryos (e). The animals had a reduced number of offspring and showed large unoccupied space inside. The gonad, although not fully visible in this image, appeared morphologically intact. Scale bar, 100 nm.

We then further aimed to evaluate our quantitative method for the determination of ribosome number in electron tomograms of animals with a reduction of RPOA-1. This protein is a DNA-directed RNA polymerase I subunit involved in the transcription of the structural RNAs for the subunits of the ribosomes.^36,37^ We hypothesized that a RNAi-mediated reduction of this subunit should lead to a compromised DNA binding and a reduction of the DNA-directed 5’-3’ RNA polymerase activity. This should lead to a statistically significant reduction of ribosomes in all tissues of the worm. For this, we prepared *rpoa-1(RNAi)* worms for electron tomography by high-pressure freezing, freeze-substitution and resin-embedding. Then we acquired electron tomograms of both germ cells and somatic cells at similar positions as analyzed in control worms. A lower number of ribosomes in these worms was already obvious by visual inspection of selected tomographic slices of the chosen different cell types in the *rpoa-1(RNAi)* worms (Fig. 4E-G). Our quantitative analysis revealed that distal cells of the gonad contained on average 17,388 ± 6,352 ribosomes per μm³, whereas cells in pachytene showed 18,668 ± 5,847 and somatic cells of the vulva 16,043 ± 9,565 ribosomes per 1 μm³, respectively. Using Welch’s t-test, we observed a significant higher number of ribosomes in control compared to *rpoa-1(RNAi)* worms in the distal part of the gonad (p = 9.8 * 10^-9^) and in the pachytene region (p = 0.004587). However, the distributions between control and *rpoa-1(RNAi)* worms in somatic vulva cells seemed not to be changed (p = 0.4937) (Fig. 4H). Worms treated in this way, also showed a reduction in offspring and exhibited large, unoccupied spaces inside (Fig. 4I). However, the gonad was superficially still intact and the vulva seemed structurally not to be affected by this treatment.

This experiment clearly demonstrated that differences in ribosome number can also be detected in cells of different origin in both control and RNAi-treated worms.

## Discussion

Here we developed a new method for the quantification of ribosome number in electron tomograms taken from specimens after high-pressure freezing, freeze substitution and resin-embedding. We demonstrated that our new approach can be used for ribosome quantification in different cell types as well as in different model systems.

### Methodological considerations

For the acquisition of tomograms from hTERT-RPE-1 cells, some critical steps during sample preparation had to be considered. 1) We avoided a trypsin-treatment during cell harvesting for high-pressure freezing as cells treated with trypsin-EDTA exhibited ribosome clustering.^38^ Instead, we applied the shake-off approach to detach cells from the support with the intention to minimize artifacts due to the use of chemicals.^39,40^ 2) Also relevant to the *C. elegans* samples, the electron beam causes a collapse of the sections in the electron microscope during data acquisition.^41^ We calculated a mean shrinkage factor to be applied to the Z-voxel size to correct for this collapse in the Z-dimension and hence to correct the volumetric measurements. Importantly, a shrinkage factor also has to be applied for different resins used for 3D electron microscopy of plastic sections as there are known variations in section shrinkage due to exposure of the electron beam.^11,42^ As we routinely only use Epon/Araldite as a resin for electron tomography, we could make use of a huge collection of samples in our lab to calculate a robust mean shrinkage factor, specific for this resin (see Suppl. Data 5). This was important as we observed quite a huge variation in section thickness in our investigated samples. This could happen during ultramicrotomy and usually averages out if serial sections were stitched together to larger volumes. However, as we relied here on volumetric measurements of single sections we thought applying a mean shrinkage factor would be the most precise way to determine the ribosome number per given volume. 3) We applied confocal light microscopy to determine both the volume of the cells and the nuclei in the hTERT-RPE-1 cells. Simply subtracting the volume of the nuclei from the total volume gave us an estimate about the whole cytoplasmic volume in hTERT-RPE-1 cells. However, the cytoplasmic volume available for ribosome distribution should be actually lower because non-ribosomal compartments such as mitochondria, Golgi, ER and densely-packed cytoskeletal regions such as F-actin also occupy space within cells. For simplicity, these additional organelles were not considered here and therefore our tomography-based calculations might actually overestimate the total number of ribosomes per cell because of this. In addition, we selected small tomographic sub-volumes in areas which did not show any organelles at all. Thus, there are a number of possibilities to fine-tune our microscopic approach for a quantitative analysis of the total number of ribosomes in whole cells. In general, a much larger number of tomograms should be analyzed in the future to get more precise information about total numbers of ribosomes in different cell types. Nevertheless, our electron tomography data showed consistent ribosome counts across different hTERT-RPE-1 cells, confirming the overall reliability of our newly developed approach.

### Quantification of ribosomes in hTERT-RPE-1 using capillary electrophoresis

Various methods can be applied for ribosome quantification.^8^ One of them is capillary electrophoresis, a routinely applied method to measure rRNA concentrations.^43^ Capillary electrophoresis is commonly used for RNA analysis due to the migration of negatively charged RNAs toward the anode in an electric field.^44^ This method, however, comes with inaccuracies related to the sample and the threshold or baseline adjustments of the RNA peaks. In our experiments, the biochemical method yielded approximately half of the ribosome count compared to the electron tomography image-analysis approach. An incomplete estimation of the non-ribosome volumes as discussed above may contribute to this discrepancy. However, the average RNA concentration from two million cells was approximately double that from one million cells (see Fig. 3C). The quantification of ribosomes in hTERT-RPE-1 cells using capillary electrophoresis provided detailed insights into rRNA content, allowing a validation of electron tomography as a new quantification method. Biochemical experiments were performed on cells harvested before the fifth passage to avoid the selection of lab derived phenotypes possibly affecting ribosome numbers as reported for Caco-2 cells.^45^ Using capillary electrophoresis, the quality of RNA samples was verified through the RNA integrity number (RIN) and the 18S/28S rRNA ratio. High RIN values (>9.5) and appropriate rRNA ratios (1.9:1 to 2:1) indicated non-degraded RNA, ensuring accurate ribosome quantification.^31,46–48^ Variations in total RNA concentrations between samples may be due to differences in sample acquisition and RNA isolation.^49^ Despite this, consistent numbers of 18S and 28S particles were observed across different sample concentrations, reflecting the co-transcription of these subunits ^50^. Overall, the biochemical measurement rRNA is relying on many factors and might always underestimate the total amount of ribosomes, as in every step of purification there is some loss of RNA and of course there is always the risk of RNA degradation. Hence, it is not very surprising that our biochemical estimation of ribosome number is lower than what we detected directly with our electron tomography image-analysis approach. As both values are only two-fold different, we speculate that the real number of ribosomes in hTERT-RPE-1 cells is in between the two values measured with both techniques.

### Tomographic quantification of ribosomes in C. elegans samples

It was previously shown that interfering with one component of RNA polymerase I disrupts transcriptional activity, thereby affecting ribosome synthesis.^51^ We aimed to analyze a difference in ribosome level by electron tomography. We used both control and *rpoa-1(RNAi)* animals to determine and compare the number of ribosomes in defined tomographic volumes, measured as ribosomes per µm^3^. Previous work has shown that a RPOA-2 RNA polymerase I B subunit mutation in *C. elegans* induced cell apoptosis and decreased ribosomal proteins to 70% compared to wild type. This decrease in ribosome level was reported together with an apoptotic phenotype of the RNA polymerase I mutant.^52^ These findings were further supported by showing that cells with defective ribosomes or low ribosome levels undergo a regulatory mechanism leading to cell death. Taking these previous findings into account, we expected control worms to exhibit a higher number of ribosomes compared to *rpoa-1(RNAi)* worms for all analyzed germ cells and somatic cells. Indeed, the RNAi-treated worms showed significant lower numbers of ribosomes in the two areas of the gonad that we investigated. However, the somatic cells in the vulva seemed not to be effected in a similar way. We speculate that the gonad as reproductive organ is more prone to this regulatory effect on the ribosome number or more sensitive to RNA-mediated interference in general compared to the somatic cells of the vulva. Another assumption could be that the low number of cell divisions and reduced protein synthesis in some somatic tissues could lead to a reduced effect. It could also mean that the number of samples investigated were too low to showcase smaller effects of ribosome downregulation.

Taken together, our newly developed approach using electron tomography can be used for a quantitative analysis of ribosome number in different cell types, in the context of complex tissues and/or different model systems. This can open up new possibilities for analyzing ribosome number or distribution by combining biochemical approaches with 3D electron microscopy.

## Acknowledgments

The authors are grateful to Silke Tulok, Dr. Michael Gerlach and Dr. Anja Nobst (Core Facility Cellular Imaging at the Carl Gustav Carus Faculty of Medicine at TU Dresden) for technical support in light microscopy and bioimage data management in OMERO, Chukwuebuka William Okafornta for help in handling of worms and Dr. Tobias Fürstenhaupt (MPI-CBG, Dresden) for technical assistance in electron tomography. M.E.H would like to thank Dr. Xingbo Yang (POL, TU Dresden) for advice and encouragements and Dr. Christian Massino (POL, TU Dresden) for the Inkscape tutorial and introduction in R studio. Research in the Müller-Reichert lab is supported by grants given by the Deutsche Forschungsgemeinschaft (MU1423/8-3, project no. 258577783 and MU1423/10-3, project no. 282354882) to T.M.R.

## Notes

### Competing Interest Statement

The authors have declared no competing interest.

## References

1. Traub, P., & Nomura, M. (1968). Structure and function of E. coli ribosomes. V. Reconstitution of functionally active 30S ribosomal particles from RNA and proteins. Proceedings of the National Academy of Sciences, 59(3), 777–784. 10.1073/pnas.59.3.777

2. Moraleva, A. A. (n.d.). Eukaryotic Ribosome Biogenesis: The 60S Subunit. Acta Naturae. 10.32607/actanaturae.11541

3. Moraleva, A. A., Deryabin, A. S., Rubtsov, Yu. P., Rubtsova, M. P., & Dontsova, O. A. (2022). Eukaryotic Ribosome Biogenesis: The 40S Subunit. Acta Naturae, 14(1), 14–30. 10.32607/actanaturae.11540

4. Ben-Shem, A., Garreau de Loubresse, N., Melnikov, S., Jenner, L., Yusupova, G., & Yusupov, M. (2011). The Structure of the Eukaryotic Ribosome at 3.0 Å Resolution. Science, 334(6062), 1524–1529. 10.1126/science.1212642

5. Hardiman, T., Ewald, J. C., Lemuth, K., Reuss, M., & Siemann-Herzberg, M. (2008). Quantification of rRNA in *Escherichia coli* using capillary gel electrophoresis with laser-induced fluorescence detection. Analytical Biochemistry, 374(1), 79–86. 10.1016/j.ab.2007.09.032

6. Ghaemmaghami, S., Huh, W.-K., Bower, K., Howson, R. W., Belle, A., Dephoure, N., O’Shea, E. K., & Weissman, J. S. (2003). Global analysis of protein expression in yeast. Nature, 425(6959), 737–741. 10.1038/nature02046

7. Newman, J. R. S., Ghaemmaghami, S., Ihmels, J., Breslow, D. K., Noble, M., DeRisi, J. L., & Weissman, J. S. (2006). Single-cell proteomic analysis of S. cerevisiae reveals the architecture of biological noise. Nature, 441(7095), 840–846. 10.1038/nature04785

8. Petelski, A. A., & Slavov, N. (2020). Analyzing Ribosome Remodeling in Health and Disease. PROTEOMICS, 20(17–18), 2000039. 10.1002/pmic.202000039

9. Winter, E. S., Schwarz, A., Fabig, G., Feldman, J. L., Pires-daSilva, A., Müller-Reichert, T., Sadler, P. L., & Shakes, D. C. (2017). Cytoskeletal variations in an asymmetric cell division support diversity in nematode sperm size and sex ratios. *Development (Cambridge*, England), 144(18), 3253–3263. 10.1242/dev.153841

10. Fabig, G., Kiewisz, R., Lindow, N., Powers, J. A., Cota, V., Quintanilla, L. J., Brugués, J., Prohaska, S., Chu, D. S., & Müller-Reichert, T. (2020). Male meiotic spindle features that efficiently segregate paired and lagging chromosomes. eLife, 9, e50988. 10.7554/eLife.50988

11. Kiewisz, R., Fabig, G., Conway, W., Baum, D., Needleman, D., & Müller-Reichert, T. (2022). Three-dimensional structure of kinetochore-fibers in human mitotic spindles. eLife, 11, e75459. 10.7554/eLife.75459

12. Wright, R. (2000). Transmission electron microscopy of yeast. Microscopy Research and Technique, 51(6), 496–510. 10.1002/1097-0029(20001215)51:6<496::AID-JEMT2>3.0.CO;2-9

13. Viotti, C., Krüger, F., Krebs, M., Neubert, C., Fink, F., Lupanga, U., Scheuring, D., Boutté, Y., Frescatada-Rosa, M., Wolfenstetter, S., Sauer, N., Hillmer, S., Grebe, M., & Schumacher, K. (2013). The endoplasmic reticulum is the main membrane source for biogenesis of the lytic vacuole in Arabidopsis. The Plant Cell, 25(9), 3434–3449. 10.1105/tpc.113.114827

14. Structome analysis of Escherichia coli cells by serial ultrathin sectioning reveals the precise cell profiles and the ribosome density | Microscopy | Oxford Academic. (n.d.). Retrieved September 6, 2024, from https://academic.oup.com/jmicro/article/66/4/283/3867840

15. Kiewisz, R., Baum, D., Müller-Reichert, T., & Fabig, G. (2023). Serial-section Electron Tomography and Quantitative Analysis of Microtubule Organization in 3D-reconstructed Mitotic Spindles. Bio-Protocol, 13(20), e4849. 10.21769/BioProtoc.4849

16. Kiewisz, R., Müller-Reichert, T., & Fabig, G. (2021). Chapter 7—High-throughput screening of mitotic mammalian cells for electron microscopy using classic histological dyes. In T. Müller-Reichert & P. Verkade (Eds.), Methods in Cell Biology (Vol. 162, pp. 151–170). Academic Press. 10.1016/bs.mcb.2020.08.005

17. Fabig, G., Schwarz, A., Striese, C., Laue, M., & Müller-Reichert, T. (2019). *In situ* analysis of male meiosis in *C. elegans*. In T. Müller-Reichert & G. Pigino (Eds.), Methods in Cell Biology (Vol. 152, pp. 119–134). Academic Press. 10.1016/bs.mcb.2019.03.013

18. Müller-Reichert, T., Hohenberg, H., O’Toole, E. T., & Mcdonald, K. (2003). Cryoimmobilization and three-dimensional visualization of C. elegans ultrastructure. Journal of Microscopy, 212(1), 71–80. 10.1046/j.1365-2818.2003.01250.x

19. Mastronarde, D. N. (2003). SerialEM: A Program for Automated Tilt Series Acquisition on Tecnai Microscopes Using Prediction of Specimen Position. Microscopy and Microanalysis, 9(S02), 1182–1183. 10.1017/S1431927603445911

20. Mastronarde, D. N. (2005). Automated electron microscope tomography using robust prediction of specimen movements. Journal of Structural Biology, 152(1), 36–51. 10.1016/j.jsb.2005.07.007

21. Mastronarde, D. N. (1997). Dual-Axis Tomography: An Approach with Alignment Methods That Preserve Resolution. Journal of Structural Biology, 120(3), 343–352. 10.1006/jsbi.1997.3919

22. Kremer, J. R., Mastronarde, D. N., & McIntosh, J. R. (1996). Computer Visualization of Three-Dimensional Image Data Using IMOD. Journal of Structural Biology, 116(1), 71–76. 10.1006/jsbi.1996.0013

23. Mastronarde, D. N., & Held, S. R. (2017). Automated tilt series alignment and tomographic reconstruction in IMOD. Journal of Structural Biology, 197(2), 102–113. 10.1016/j.jsb.2016.07.011

24. Schindelin, J., Arganda-Carreras, I., Frise, E., Kaynig, V., Longair, M., Pietzsch, T., Preibisch, S., Rueden, C., Saalfeld, S., Schmid, B., Tinevez, J.-Y., White, D. J., Hartenstein, V., Eliceiri, K., Tomancak, P., & Cardona, A. (2012). Fiji: An open-source platform for biological-image analysis cvb. Nature Methods, 9(7), 676–682. 10.1038/nmeth.2019

25. Gusnard, D., & Kirschner, R. H. (1977). Cell and organelle shrinkage during preparation for scanning electron microscopy: Effects of fixation, dehydration and critical point drying. Journal of Microscopy, 110(1), 51–57. 10.1111/j.1365-2818.1977.tb00012.x

26. Luther, P. K. (2006). Sample Shrinkage and Radiation Damage of Plastic Sections. In J. Frank (Ed.), Electron Tomography: Methods for Three-Dimensional Visualization of Structures in the Cell (pp. 17–48). Springer. 10.1007/978-0-387-69008-7_2

27. Yi, X., Verbeke, E. J., Chang, Y., Dickinson, D. J., & Taylor, D. W. (2019). Electron microscopy snapshots of single particles from single cells. The Journal of Biological Chemistry, 294(5), 1602–1608. 10.1074/jbc.RA118.006686

28. Cox, R. A., & Arnstein, H. R. V. (2003). Translation of RNA to Protein. In R. A. Meyers (Ed.), Encyclopedia of Physical Science and Technology (Third Edition) (pp. 31–51). Academic Press. 10.1016/B0-12-227410-5/00788-2

29. Chomczynski, P., & Sacchi, N. (1987). Single-step method of RNA isolation by acid guanidinium thiocyanate-phenol-chloroform extraction. Analytical Biochemistry, 162(1), 156–159. 10.1016/0003-2697(87)90021-2

30. Nachamkin, I., Panaro, N. J., Li, M., Ung, H., Yuen, P. K., Kricka, L. J., & Wilding, P. (2001). Agilent 2100 Bioanalyzer for Restriction Fragment Length Polymorphism Analysis of the Campylobacter jejuni Flagellin Gene. Journal of Clinical Microbiology, 39(2), 754–757. 10.1128/jcm.39.2.754-757.2001

31. Schroeder, A., Mueller, O., Stocker, S., Salowsky, R., Leiber, M., Gassmann, M., Lightfoot, S., Menzel, W., Granzow, M., & Ragg, T. (2006). The RIN: An RNA integrity number for assigning integrity values to RNA measurements. BMC Molecular Biology, 7, 3. 10.1186/1471-2199-7-3

32. Brenner, S. (1974). THE GENETICS OF CAENORHABDITIS ELEGANS. Genetics, 77(1), 71–94. 10.1093/genetics/77.1.71

33. Hammell, C. M., & Hannon, G. J. (2012). Inducing RNAi in C. elegans by Feeding with dsRNA-expressing E. coli. Cold Spring Harbor Protocols, 2012(12), pdb.prot072348. 10.1101/pdb.prot072348

34. Fire, A., Xu, S., Montgomery, M. K., Kostas, S. A., Driver, S. E., & Mello, C. C. (1998). Potent and specific genetic interference by double-stranded RNA in Caenorhabditis elegans. Nature, 391(6669), 806–811. 10.1038/35888

35. Kamath, R. S., & Ahringer, J. (2003). Genome-wide RNAi screening in *Caenorhabditis elegans*. Methods, 30(4), 313–321. 10.1016/S1046-2023(03)00050-1

36. Kane, C. (2013). Transcription. In S. Maloy & K. Hughes (Eds.), Brenner’s Encyclopedia of Genetics (Second Edition) (pp. 85–92). Academic Press. 10.1016/B978-0-12-374984-0.01551-5

37. Pu, Y.-Z., Wan, Q.-L., Ding, A.-J., Luo, H.-R., & Wu, G.-S. (2017). Quantitative proteomics analysis of *Caenorhabditis elegans* upon germ cell loss. Journal of Proteomics, 156, 85–93. 10.1016/j.jprot.2017.01.011

38. Furcht, L. T., & Wendelschafer-Crabb, G. (1978). Trypsin-induced coordinate alterations in cell shape, cytoskeleton, and intrinsic membrane structure of contact-inhibited cells. Experimental Cell Research, 114(1), 1–14. 10.1016/0014-4827(78)90029-0

39. Fox, M. H., Read, R. A., & Bedford, J. S. (1987). Comparison of synchronized Chinese hamster ovary cells obtained by mitotic shake-off, hydroxyurea, aphidicolin, or methotrexate. Cytometry, 8(3), 315–320. 10.1002/cyto.990080312

40. Terasima, T., & Tolmach, L. J. (1963). Growth and nucleic acid synthesis in synchronously dividing populations of HeLa cells. Experimental Cell Research, 30(2), 344–362. 10.1016/0014-4827(63)90306-9

41. Luther, P. K., Lawrence, M. C., & Crowther, R. A. (1988). A method for monitoring the collapse of plastic sections as a function of electron dose. Ultramicroscopy, 24(1), 7–18. 10.1016/0304-3991(88)90322-1

42. Kizilyaprak, C., Longo, G., Daraspe, J., & Humbel, B. M. (2015). Investigation of resins suitable for the preparation of biological sample for 3-D electron microscopy. Journal of Structural Biology, 189(2), 135–146. 10.1016/j.jsb.2014.10.009

43. Pfaffl, M. W., Fleige, S., & Riedmaier, I. (2008). Validation of Lab-on-Chip Capillary Electrophoresis Systems for Total RNA Quality and Quantity Control. Biotechnology & Biotechnological Equipment, 22(3), 829–834. 10.1080/13102818.2008.10817562

44. Rio, D. C., Ares, M., Hannon, G. J., & Nilsen, T. W. (2010). Nondenaturing Agarose Gel Electrophoresis of RNA. Cold Spring Harbor Protocols, 2010(6), pdb.prot5445. 10.1101/pdb.prot5445

45. Yu, H., Cook, T. J., & Sinko, P. J. (1997). Evidence for Diminished Functional Expression of Intestinal Transporters in Caco-2 Cell Monolayers at High Passages. Pharmaceutical Research, 14(6), 757–762. 10.1023/A:1012150405949

46. Skrypina, N. A., Timofeeva, A. V., Khaspekov, G. L., Savochkina, L. P., & Beabealashvilli, R. Sh. (2003). Total RNA suitable for molecular biology analysis. Journal of Biotechnology, 105(1), 1–9. 10.1016/S0168-1656(03)00140-8

47. Imbeaud, S., Graudens, E., Boulanger, V., Barlet, X., Zaborski, P., Eveno, E., Mueller, O., Schroeder, A., & Auffray, C. (2005). Towards standardization of RNA quality assessment using user-independent classifiers of microcapillary electrophoresis traces. Nucleic Acids Research, 33(6), e56. 10.1093/nar/gni054

48. Auer, H., Lyianarachchi, S., Newsom, D., Klisovic, M. I., Marcucci, U., & Kornacker, K. (2003). Chipping away at the chip bias: RNA degradation in microarray analysis. Nature Genetics, 35(4), 292–293. 10.1038/ng1203-292

49. Tricarico, C., Pinzani, P., Bianchi, S., Paglierani, M., Distante, V., Pazzagli, M., Bustin, S. A., & Orlando, C. (2002). Quantitative real-time reverse transcription polymerase chain reaction: Normalization to rRNA or single housekeeping genes is inappropriate for human tissue biopsies. Analytical Biochemistry, 309(2), 293–300. 10.1016/S0003-2697(02)00311-1

50. Lafontaine, D. L. J., & Tollervey, D. (1998). Birth of the snoRNPs: The evolution of the modification-guide snoRNAs. Trends in Biochemical Sciences, 23(10), 383–388. 10.1016/S0968-0004(98)01260-2

51. Laferté, A., Favry, E., Sentenac, A., Riva, M., Carles, C., & Chédin, S. (2006). The transcriptional activity of RNA polymerase I is a key determinant for the level of all ribosome components. Genes & Development, 20(15), 2030–2040. 10.1101/gad.386106

52. Eberhard, R., Stergiou, L., Hofmann, E. R., Hofmann, J., Haenni, S., Teo, Y., Furger, A., & Hengartner, M. O. (2013). Ribosome Synthesis and MAPK Activity Modulate Ionizing Radiation-Induced Germ Cell Apoptosis in Caenorhabditis elegans. PLOS Genetics, 9(11), e1003943. 10.1371/journal.pgen.1003943

