## Supplementary material for "*In situ* quantification of ribosome number by electron tomography": Suppl. Data 6

Assay Class: Eukaryote Total RNA Nano
Data Path: C:\...Eukaryote Total RNA Nano_DEDAE01485_2022-06-14_15-36-23.xad

Created: 6/14/2022 3:36:23 PM
Modified: 6/14/2022 4:08:38 PM

### Electrophoresis File Run Summary

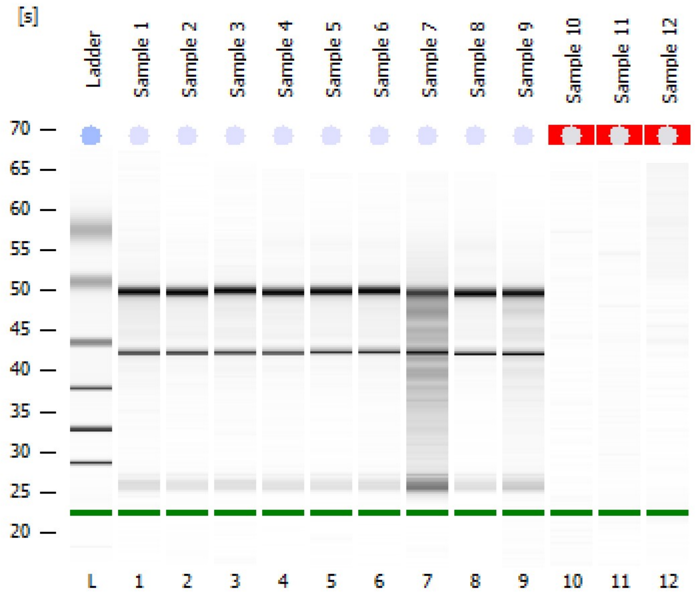


Instrument Information:

| | | | |
|---|---|---|---|
| Instrument Name: | DEDAE01485 | Firmware: | C.01.069 |
| Serial#: | DEDAE01485 | Type: | G2939B |

Assay Information:

| | |
|---|---|
| Assay Origin Path: | C:\Program Files (x86)\Agilent\2100 bioanalyzer\2100 expert\assays\RNA\Eukaryote Total RNA Nano Series II.xsy |
| Assay Class: | Eukaryote Total RNA Nano |
| Version: | 2.6 |
| Assay Comments: | Total RNA Analysis ng sensitivity (Eukaryote) |

© Copyright 2003 - 2009 Agilent Technologies, Inc.

Chip Information:

Chip Lot #:

Reagent Kit Lot #:

Chip Comments:

**Sample 1**

RIN: 9.50

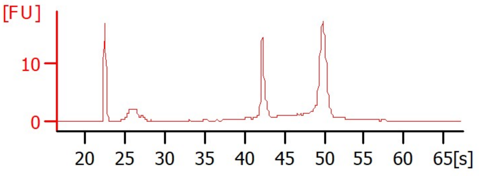

**Sample 2**

RIN: 9.80

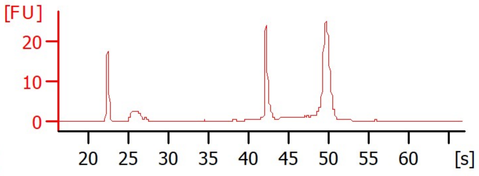

**Sample 3**

RIN: 9.80

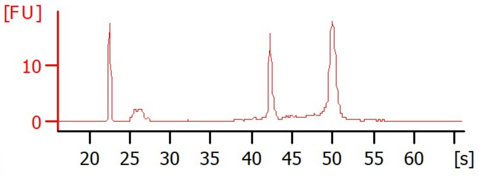

**Sample 4**

RIN: 9.90

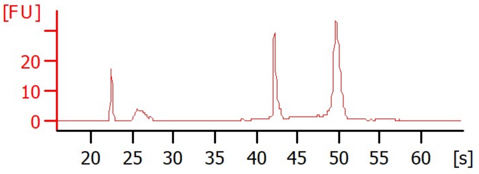

**Sample 5**

RIN:10

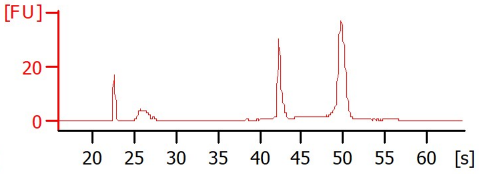

**Sample 6**

RIN: 9.80

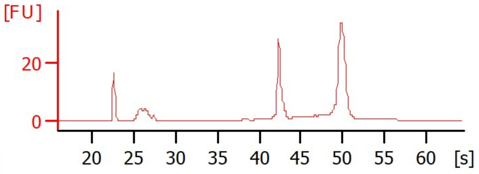

**Sample 7**

RIN: 5.80

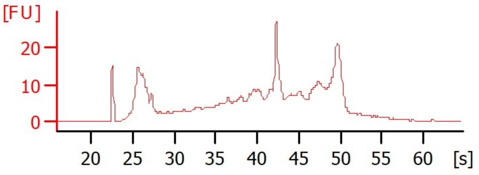

**Sample 8**

RIN: 9.70

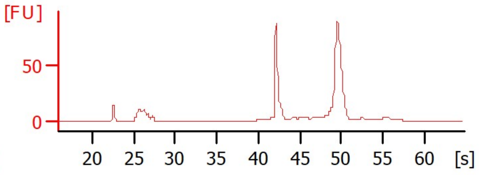

**Sample 9**

RIN: 8.40

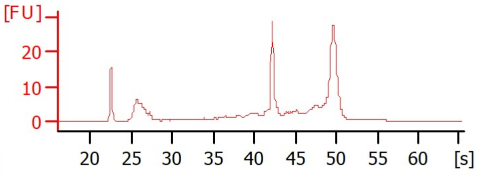

**Sample 10**

RIN N/A

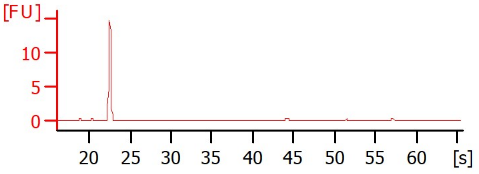

**Sample 11**

RIN N/A

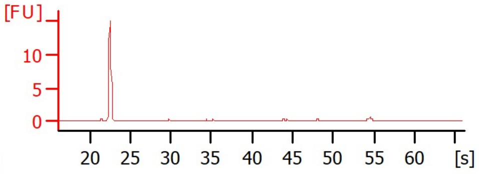

**Sample 12**

RIN N/A

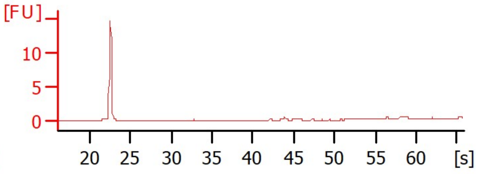

Assay Class: Eukaryote Total RNA Nano
Data Path: C:\...Eukaryote Total RNA Nano_DEDAE01485_2022-06-14_15-36-23.xad
Created: 6/14/2022 3:36:23 PM
Modified: 6/14/2022 4:08:38 PM

### Electrophoresis File Run Summary (Chip Summary)

| Sample Name | Sample Comment | Status | Result Label | Result Color |
|---|---|---|---|---|
| Sample 1 | | ✔ | RIN: 9.50 | |
| Sample 2 | | ✔ | RIN: 9.80 | |
| Sample 3 | | ✔ | RIN: 9.80 | |
| Sample 4 | | ✔ | RIN: 9.90 | |
| Sample 5 | | ✔ | RIN:10 | |
| Sample 6 | | ✔ | RIN: 9.80 | |
| Sample 7 | | ✔ | RIN: 5.80 | |
| Sample 8 | | ✔ | RIN: 9.70 | |
| Sample 9 | | ✔ | RIN: 8.40 | |
| Sample 10 | | ✔ | RIN N/A | |
| Sample 11 | | ✔ | RIN N/A | |
| Sample 12 | | ✔ | RIN N/A | |
| Ladder | | ✔ | All Other Samples | |

**Chip Lot #** **Reagent Kit Lot #**

**Chip Comments :**


© Copyright 2003 - 2018 Agilent, Inc.

Assay Class: Eukaryote Total RNA Nano
Data Path: C:\...Eukaryote Total RNA Nano_DEDAE01485_2022-06-14_15-36-23.xad
Created: 6/14/2022 3:36:23 PM
Modified: 6/14/2022 4:08:38 PM

### Electrophoresis Assay Details

#### General Analysis Settings

Number of Available Sample and Ladder Wells (Max.) : 13
Minimum Visible Range [s] : 17
Maximum Visible Range [s] : 70
Start Analysis Time Range [s] : 19
End Analysis Time Range [s] : 69
Ladder Concentration [ng/µl] : 150
Lower Marker Concentration [ng/µl] : 0
Upper Marker Concentration [ng/µl] : 0
Used Lower Marker for Quantitation
Standard Curve Fit is Logarithmic
Show Data Aligned to Lower Marker

#### Integrator Settings

Integration Start Time [s] : 19
Integration End Time [s] : 69
Slope Threshold : 0.6
Height Threshold [FU] : 0.5
Area Threshold : 0.2
Width Threshold [s] : 0.5
Baseline Plateau [s] : 6

#### Filter Settings

Filter Width [s] : 0.5
Polynomial Order : 4

#### Ladder

| Ladder Peak | Size |
|---|---|
| 1 | 25 |
| 2 | 200 |
| 3 | 500 |
| 4 | 1000 |
| 5 | 2000 |
| 6 | 4000 |

 © Copyright 2003 - 2018 Agilent, Inc. 

| | | | |
|---|---|---|---|
| Assay Class: | Eukaryote Total RNA Nano | Created: | 6/14/2022 3:36:23 PM |
| Data Path: | C:\...Eukaryote Total RNA Nano_DEDAE01485_2022-06-14_15-36-23.xad | Modified: | 6/14/2022 4:08:38 PM |

### Electropherogram Summary

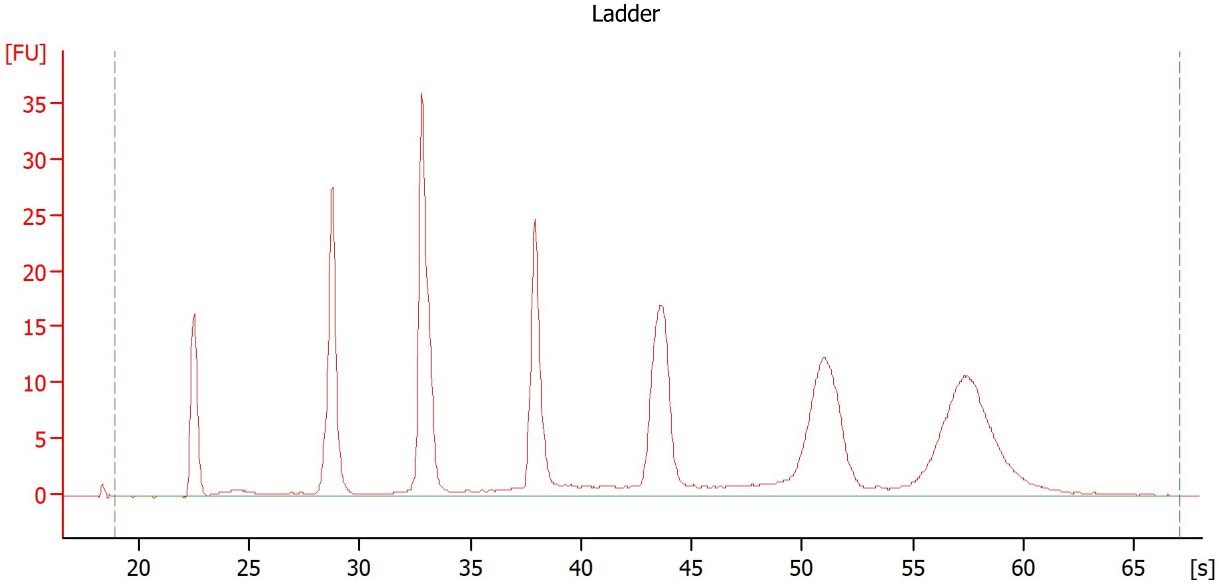


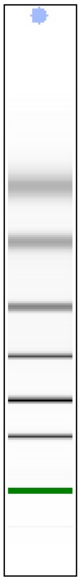

### Overall Results for Ladder

| | | | |
|---|---|---|---|
| RNA Area: | 291.0 | Result Flagging Color: | |
| RNA Concentration: | 150 ng/µl | Result Flagging Label: | All Other Samples |

 © Copyright 2003 - 2018 Agilent, Inc. 

Assay Class: Eukaryote Total RNA Nano
Data Path: C:\...Eukaryote Total RNA Nano_DEDAE01485_2022-06-14_15-36-23.xad

Created: 6/14/2022 3:36:23 PM
Modified: 6/14/2022 4:08:38 PM

**Electropherogram Summary Continued ...**

Sample 1

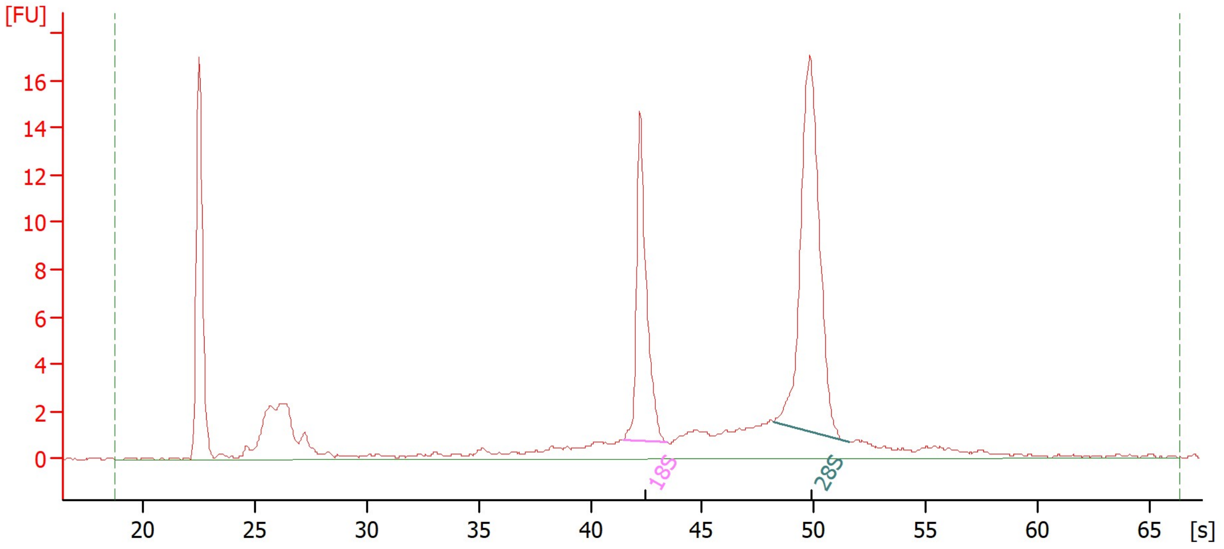


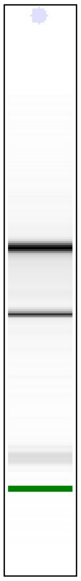

**Overall Results for sample 1 : Sample 1**

| | | | |
|---|---|---|---|
| RNA Area: | 102.8 | RNA Integrity Number (RIN): | 9.5 (B.02.10) |
| RNA Concentration: | 53 ng/µl | Result Flagging Color: | |
| rRNA Ratio [28s / 18s]: | 1.8 | Result Flagging Label: | RIN: 9.50 |

**Fragment table for sample 1 : Sample 1**

| Name | Start Time [s] | End Time [s] | Area | % of total Area |
|---|---|---|---|---|
| 18S | 41.49 | 43.48 | 16.9 | 16.5 |
| 28S | 48.17 | 51.59 | 31.1 | 30.2 |

 © Copyright 2003 - 2018 Agilent, Inc. 

| | | | |
|---|---|---|---|
| Assay Class: | Eukaryote Total RNA Nano | Created: | 6/14/2022 3:36:23 PM |
| Data Path: | C:\...Eukaryote Total RNA Nano_DEDAE01485_2022-06-14_15-36-23.xad | Modified: | 6/14/2022 4:08:38 PM |

**Electropherogram Summary Continued ...**

Sample 2

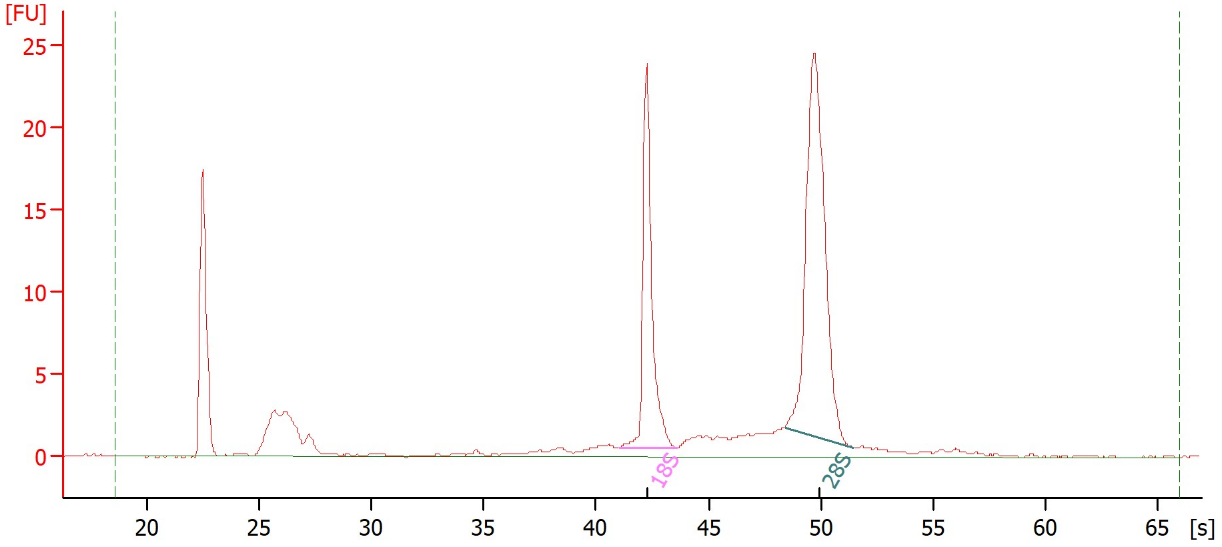


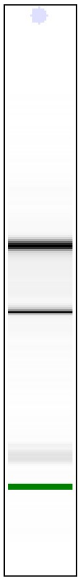

**Overall Results for sample 2 :** **Sample 2**

| | | | |
|---|---|---|---|
| RNA Area: | 122.3 | RNA Integrity Number (RIN): | 9.8 (B.02.10) |
| RNA Concentration: | 63 ng/µl | Result Flagging Color: | |
| rRNA Ratio [28s / 18s]: | 1.9 | Result Flagging Label: | RIN: 9.80 |

**Fragment table for sample 2 :** **Sample 2**

| Name | Start Time [s] | End Time [s] | Area | % of total Area |
|---|---|---|---|---|
| 18S | 40.99 | 43.61 | 24.3 | 19.8 |
| 28S | 48.39 | 51.45 | 45.2 | 37.0 |

 © Copyright 2003 - 2018 Agilent, Inc. 

| | | | |
|---|---|---|---|
| Assay Class: | Eukaryote Total RNA Nano | Created: | 6/14/2022 3:36:23 PM |
| Data Path: | C:\...Eukaryote Total RNA Nano_DEDAE01485_2022-06-14_15-36-23.xad | Modified: | 6/14/2022 4:08:38 PM |

**Electropherogram Summary Continued ...**

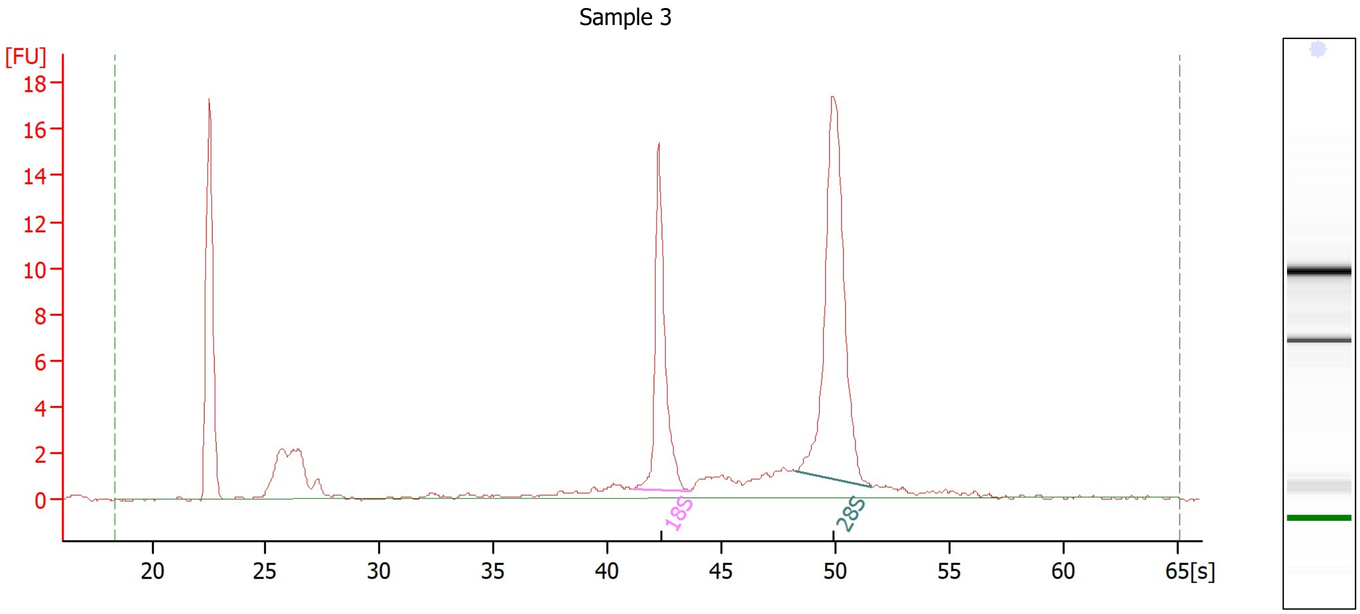


**Overall Results for sample 3 :** **Sample 3**

| | | | |
|---|---|---|---|
| RNA Area: | 90.7 | RNA Integrity Number (RIN): | 9.8 (B.02.10) |
| RNA Concentration: | 47 ng/µl | Result Flagging Color: | |
| rRNA Ratio [28s / 18s]: | 1.9 | Result Flagging Label: | RIN: 9.80 |

**Fragment table for sample 3 :** **Sample 3**

| Name | Start Time [s] | End Time [s] | Area | % of total Area |
|---|---|---|---|---|
| 18S | 41.08 | 43.63 | 16.9 | 18.6 |
| 28S | 48.25 | 51.60 | 31.8 | 35.1 |

 © Copyright 2003 - 2018 Agilent, Inc. 

| | | | |
|---|---|---|---|
| Assay Class: | Eukaryote Total RNA Nano | Created: | 6/14/2022 3:36:23 PM |
| Data Path: | C:\...Eukaryote Total RNA Nano_DEDAE01485_2022-06-14_15-36-23.xad | Modified: | 6/14/2022 4:08:38 PM |

**Electropherogram Summary Continued ...**

Sample 4

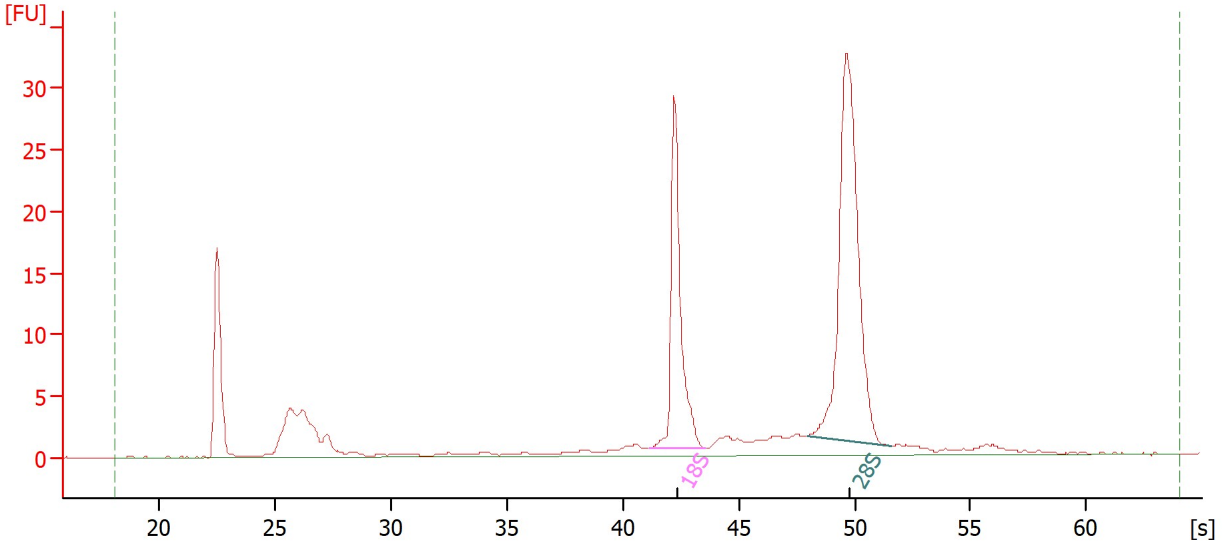


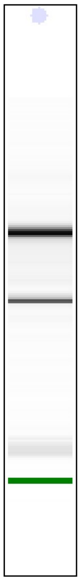

**Overall Results for sample 4 :** **Sample 4**

| | | | |
|---|---|---|---|
| RNA Area: | 157.3 | RNA Integrity Number (RIN): | 9.9 (B.02.10) |
| RNA Concentration: | 81 ng/µl | Result Flagging Color: | |
| rRNA Ratio [28s / 18s]: | 1.9 | Result Flagging Label: | RIN: 9.90 |

**Fragment table for sample 4 :** **Sample 4**

| Name | Start Time [s] | End Time [s] | Area | % of total Area |
|---|---|---|---|---|
| 18S | 41.10 | 43.52 | 31.0 | 19.7 |
| 28S | 47.97 | 51.59 | 59.9 | 38.0 |


© Copyright 2003 - 2018 Agilent, Inc.

Assay Class: Eukaryote Total RNA Nano
Data Path: C:\...Eukaryote Total RNA Nano_DEDAE01485_2022-06-14_15-36-23.xad

Created: 6/14/2022 3:36:23 PM
Modified: 6/14/2022 4:08:38 PM

**Electropherogram Summary Continued ...**

Sample 5

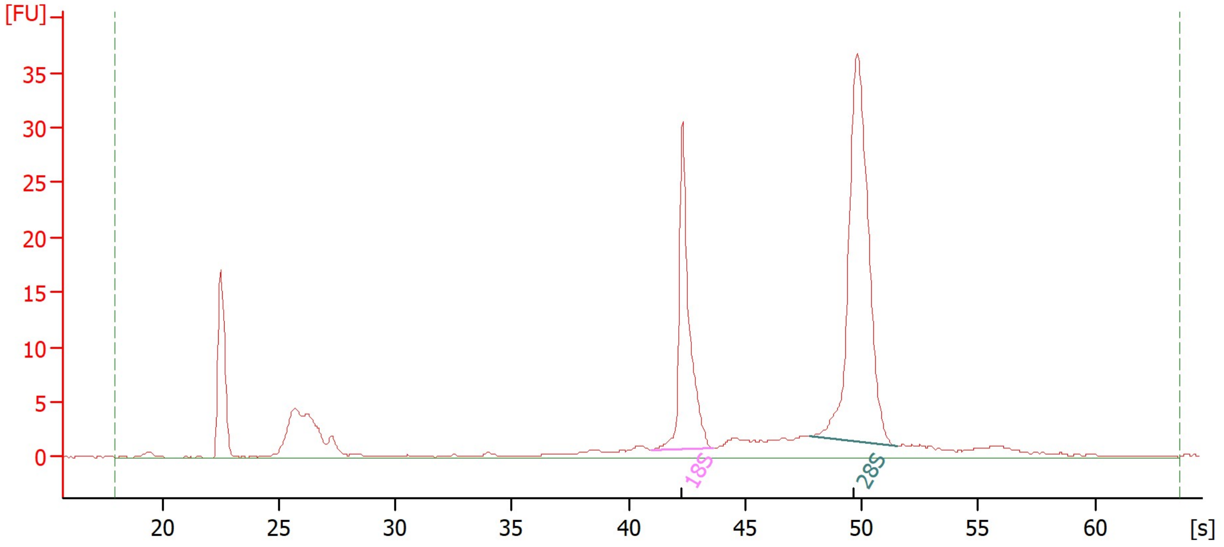


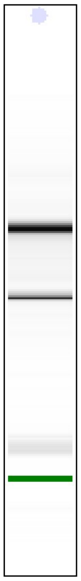

**Overall Results for sample 5 :** **Sample 5**

| | | | |
|---|---|---|---|
| RNA Area: | 182.1 | RNA Integrity Number (RIN): | 10 (B.02.10) |
| RNA Concentration: | 94 ng/µl | Result Flagging Color: | |
| rRNA Ratio [28s / 18s]: | 2.0 | Result Flagging Label: | RIN:10 |

**Fragment table for sample 5 :** **Sample 5**

| Name | Start Time [s] | End Time [s] | Area | % of total Area |
|---|---|---|---|---|
| 18S | 40.99 | 43.66 | 34.9 | 19.1 |
| 28S | 47.79 | 51.55 | 69.6 | 38.2 |

 © Copyright 2003 - 2018 Agilent, Inc. 

| | | | |
|---|---|---|---|
| Assay Class: | Eukaryote Total RNA Nano | Created: | 6/14/2022 3:36:23 PM |
| Data Path: | C:\...Eukaryote Total RNA Nano_DEDAE01485_2022-06-14_15-36-23.xad | Modified: | 6/14/2022 4:08:38 PM |

**Electropherogram Summary Continued ...**

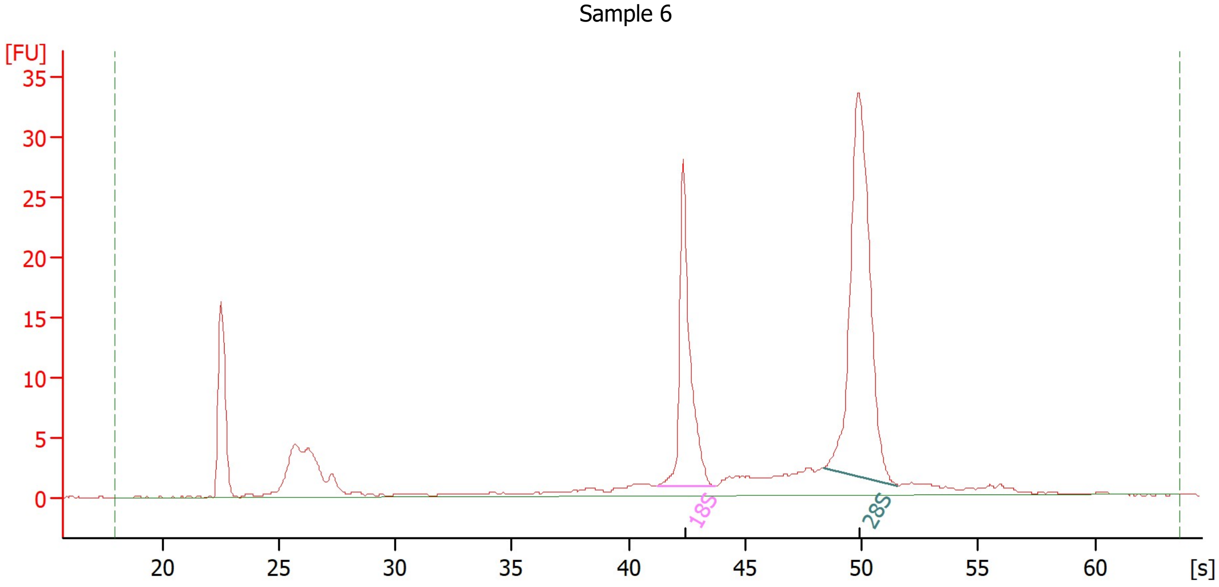


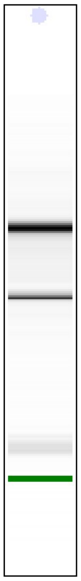

**Overall Results for sample 6 : Sample 6**

| | | | |
|---|---|---|---|
| RNA Area: | 177.7 | RNA Integrity Number (RIN): | 9.8 (B.02.10) |
| RNA Concentration: | 92 ng/µl | Result Flagging Color: | |
| rRNA Ratio [28s / 18s]: | 1.9 | Result Flagging Label: | RIN: 9.80 |

**Fragment table for sample 6 : Sample 6**

| Name | Start Time [s] | End Time [s] | Area | % of total Area |
|---|---|---|---|---|
| 18S | 41.22 | 43.76 | 32.6 | 18.4 |
| 28S | 48.37 | 51.55 | 62.3 | 35.0 |

 © Copyright 2003 - 2018 Agilent, Inc. 

| | | | |
|---|---|---|---|
| Assay Class: | Eukaryote Total RNA Nano | Created: | 6/14/2022 3:36:23 PM |
| Data Path: | C:\...Eukaryote Total RNA Nano_DEDAE01485_2022-06-14_15-36-23.xad | Modified: | 6/14/2022 4:08:38 PM |

**Electropherogram Summary Continued ...**

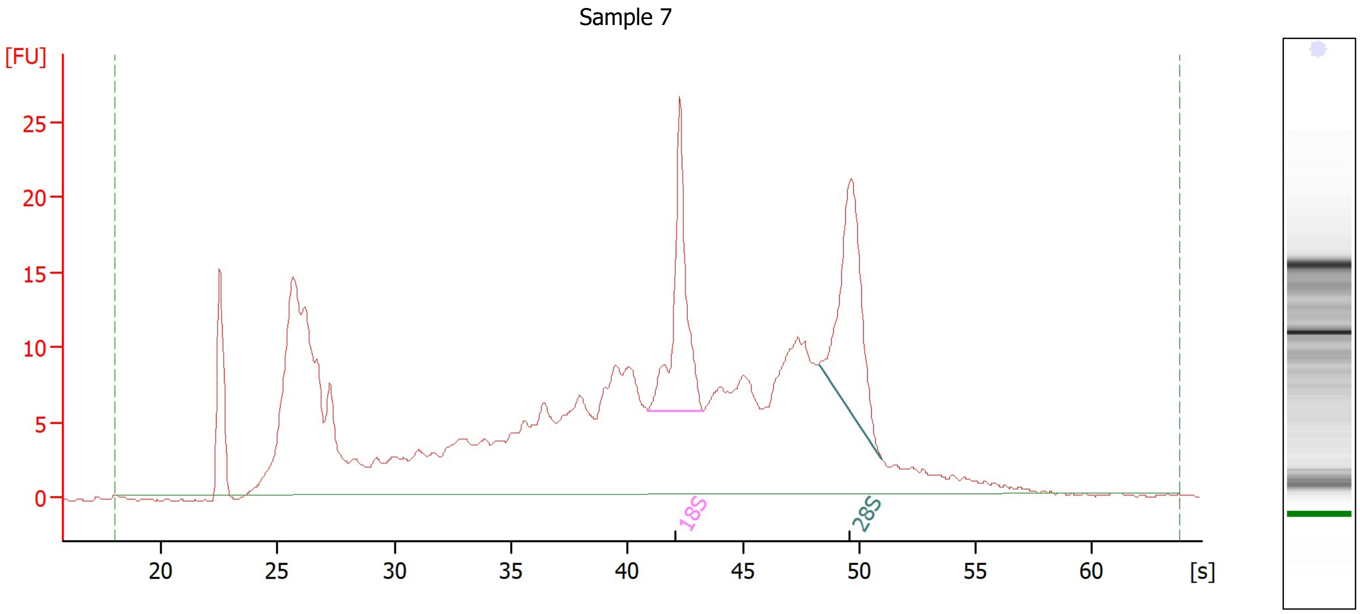


**Overall Results for sample 7 :** **Sample 7**

| | | | |
|---|---|---|---|
| RNA Area: | 476.2 | RNA Integrity Number (RIN): | 5.8 (B.02.10) |
| RNA Concentration: | 245 ng/µl | Result Flagging Color: | |
| rRNA Ratio [28s / 18s]: | 1.1 | Result Flagging Label: | RIN: 5.80 |

**Fragment table for sample 7 :** **Sample 7**

| Name | Start Time [s] | End Time [s] | Area | % of total Area |
|---|---|---|---|---|
| 18S | 40.89 | 43.29 | 28.9 | 6.1 |
| 28S | 48.28 | 50.96 | 32.8 | 6.9 |

 © Copyright 2003 - 2018 Agilent, Inc. 

| | | | |
|---|---|---|---|
| Assay Class: | Eukaryote Total RNA Nano | Created: | 6/14/2022 3:36:23 PM |
| Data Path: | C:\...Eukaryote Total RNA Nano_DEDAE01485_2022-06-14_15-36-23.xad | Modified: | 6/14/2022 4:08:38 PM |

**Electropherogram Summary Continued ...**

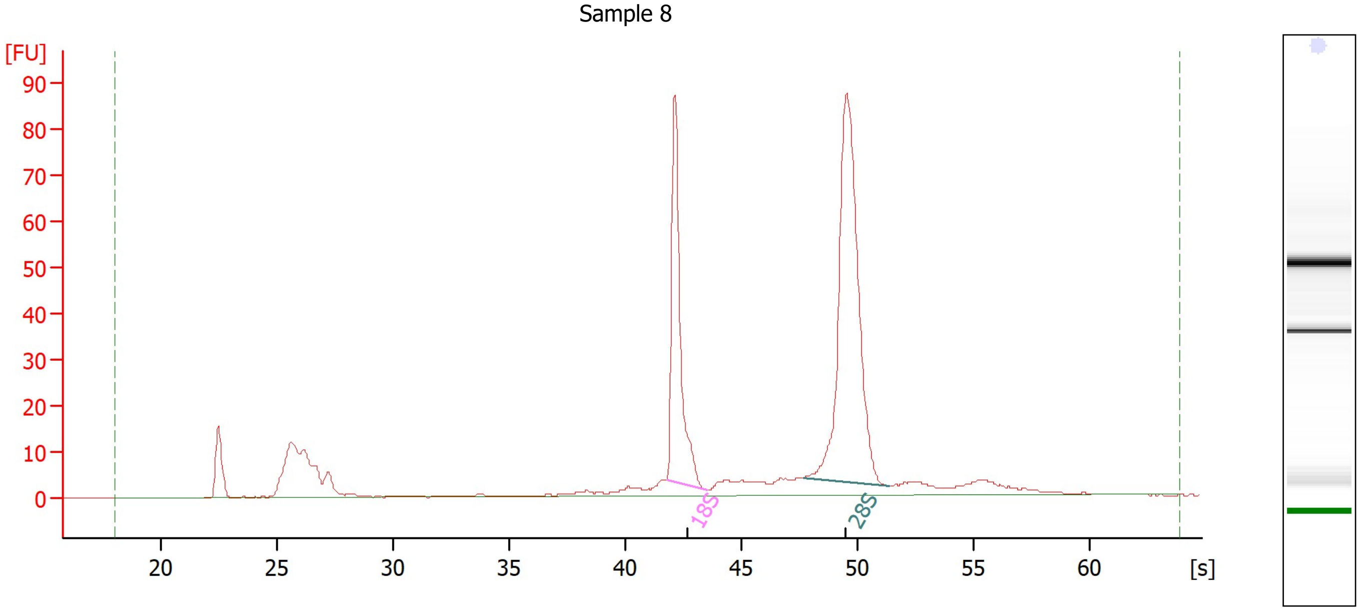


**Overall Results for sample 8 : Sample 8**

| | | | |
|---|---|---|---|
| RNA Area: | 429.3 | RNA Integrity Number (RIN): | 9.7 (B.02.10) |
| RNA Concentration: | 221 ng/µl | Result Flagging Color: | |
| rRNA Ratio [28s / 18s]: | 2.0 | Result Flagging Label: | RIN: 9.70 |

**Fragment table for sample 8 : Sample 8**

| Name | Start Time [s] | End Time [s] | Area | % of total Area |
|---|---|---|---|---|
| 18S | 41.81 | 43.52 | 82.4 | 19.2 |
| 28S | 47.69 | 51.44 | 164.1 | 38.2 |

 © Copyright 2003 - 2018 Agilent, Inc. 

Assay Class: Eukaryote Total RNA Nano
Data Path: C:\...Eukaryote Total RNA Nano_DEDAE01485_2022-06-14_15-36-23.xad

Created: 6/14/2022 3:36:23 PM
Modified: 6/14/2022 4:08:38 PM

**Electropherogram Summary Continued ...**

Sample 9

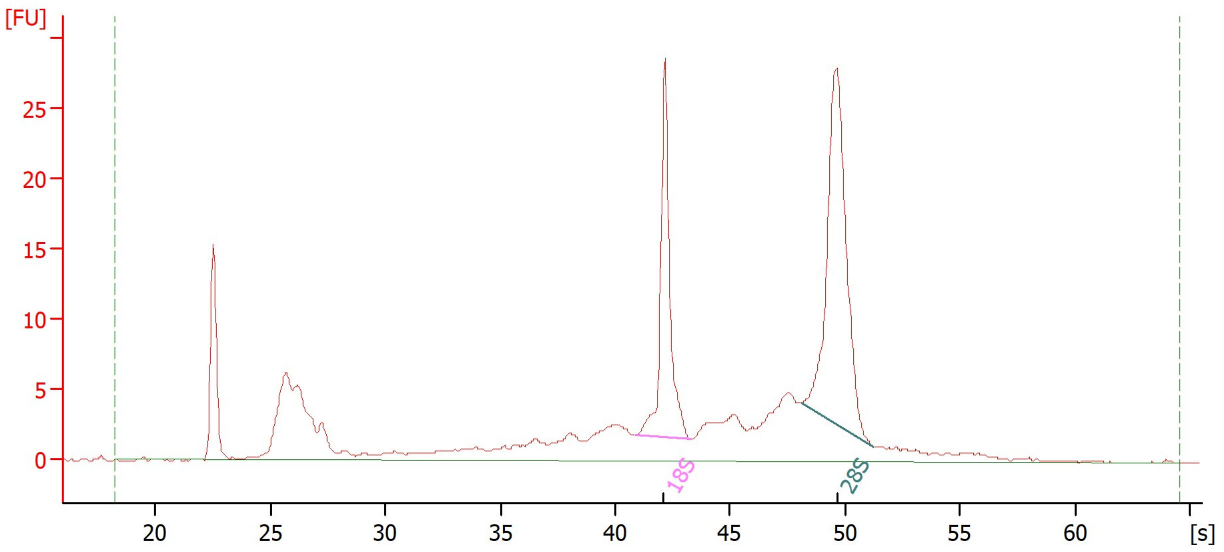


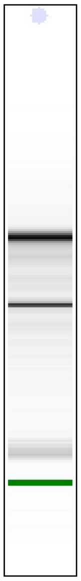

**Overall Results for sample 9 :** **Sample 9**

| | | | |
|---|---|---|---|
| RNA Area: | 219.8 | RNA Integrity Number (RIN): | 8.4 (B.02.10) |
| RNA Concentration: | 113 ng/µl | Result Flagging Color: | |
| rRNA Ratio [28s / 18s]: | 1.7 | Result Flagging Label: | RIN: 8.40 |

**Fragment table for sample 9 :** **Sample 9**

| Name | Start Time [s] | End Time [s] | Area | % of total Area |
|---|---|---|---|---|
| 18S | 40.88 | 43.32 | 29.0 | 13.2 |
| 28S | 48.09 | 51.27 | 49.4 | 22.5 |

 © Copyright 2003 - 2018 Agilent, Inc. 

| | | | |
|---|---|---|---|
| Assay Class: | Eukaryote Total RNA Nano | Created: | 6/14/2022 3:36:23 PM |
| Data Path: | C:\...Eukaryote Total RNA Nano_DEDAE01485_2022-06-14_15-36-23.xad | Modified: | 6/14/2022 4:08:38 PM |

**Electropherogram Summary Continued ...**

Sample 10

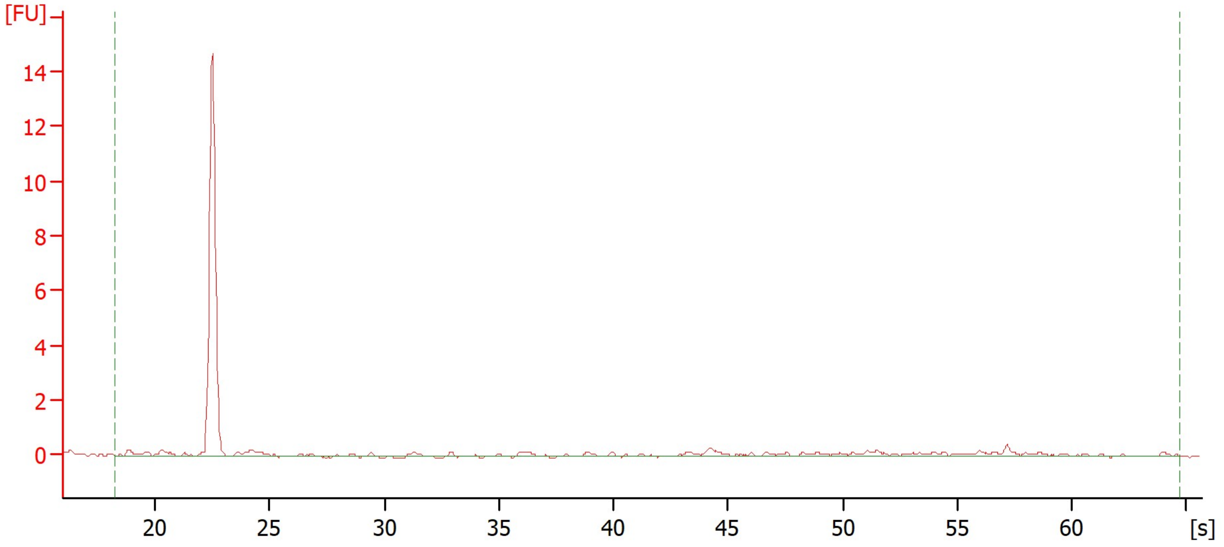


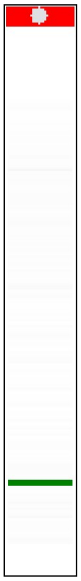

**Overall Results for sample 10 :** **<u>Sample 10</u>**

| | | | |
|---|---|---|---|
| RNA Area: | 4.6 | RNA Integrity Number (RIN): | N/A (B.02.10) |
| RNA Concentration: | 2 ng/µl | Result Flagging Color: | |
| rRNA Ratio [28s / 18s]: | 0.0 | Result Flagging Label: | RIN N/A |

 © Copyright 2003 - 2018 Agilent, Inc. 

| | | | |
|---|---|---|---|
| Assay Class: | Eukaryote Total RNA Nano | Created: | 6/14/2022 3:36:23 PM |
| Data Path: | C:\...Eukaryote Total RNA Nano_DEDAE01485_2022-06-14_15-36-23.xad | Modified: | 6/14/2022 4:08:38 PM |

**Electropherogram Summary Continued ...**

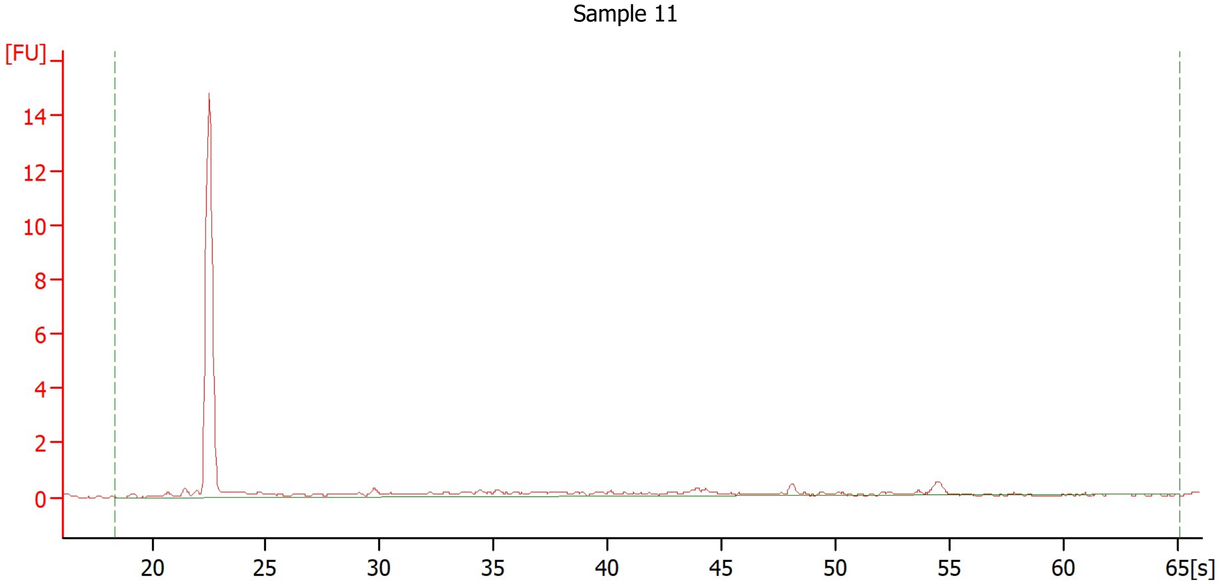


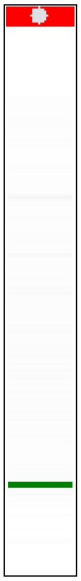

**Overall Results for sample 11 : Sample 11**

| | | | |
|---|---|---|---|
| RNA Area: | 9.8 | RNA Integrity Number (RIN): | N/A (B.02.10) |
| RNA Concentration: | 5 ng/µl | Result Flagging Color: | |
| rRNA Ratio [28s / 18s]: | 0.0 | Result Flagging Label: | RIN N/A |

 © Copyright 2003 - 2018 Agilent, Inc. 

| | | | |
|---|---|---|---|
| Assay Class: | Eukaryote Total RNA Nano | Created: | 6/14/2022 3:36:23 PM |
| Data Path: | C:\...Eukaryote Total RNA Nano_DEDAE01485_2022-06-14_15-36-23.xad | Modified: | 6/14/2022 4:08:38 PM |

**Electropherogram Summary Continued ...**

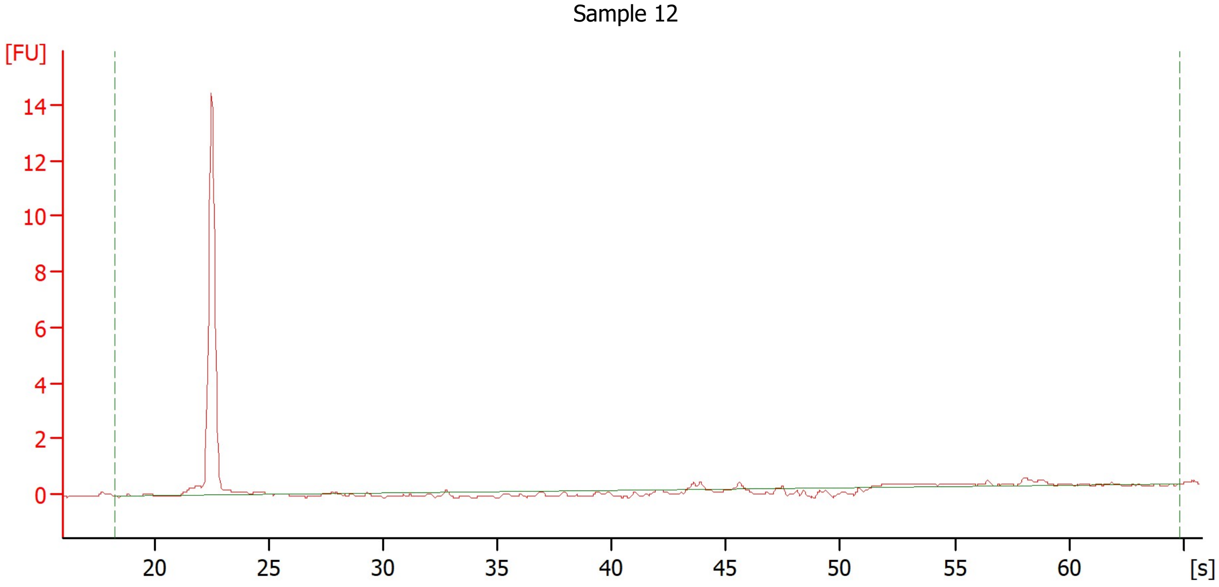


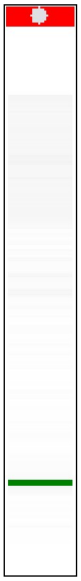

**Overall Results for sample 12 :** **Sample 12**

| | | | |
|---|---|---|---|
| RNA Area: | 3.2 | RNA Integrity Number (RIN): | N/A (B.02.10) |
| RNA Concentration: | 2 ng/µl | Result Flagging Color: | |
| rRNA Ratio [28s / 18s]: | 0.0 | Result Flagging Label: | RIN N/A |

 © Copyright 2003 - 2018 Agilent, Inc. 

Assay Class: Eukaryote Total RNA Nano
Data Path: C:\...Eukaryote Total RNA Nano_DEDAE01485_2022-06-14_15-36-23.xad

Created: 6/14/2022 3:36:23 PM
Modified: 6/14/2022 4:08:38 PM

**Gel Image**

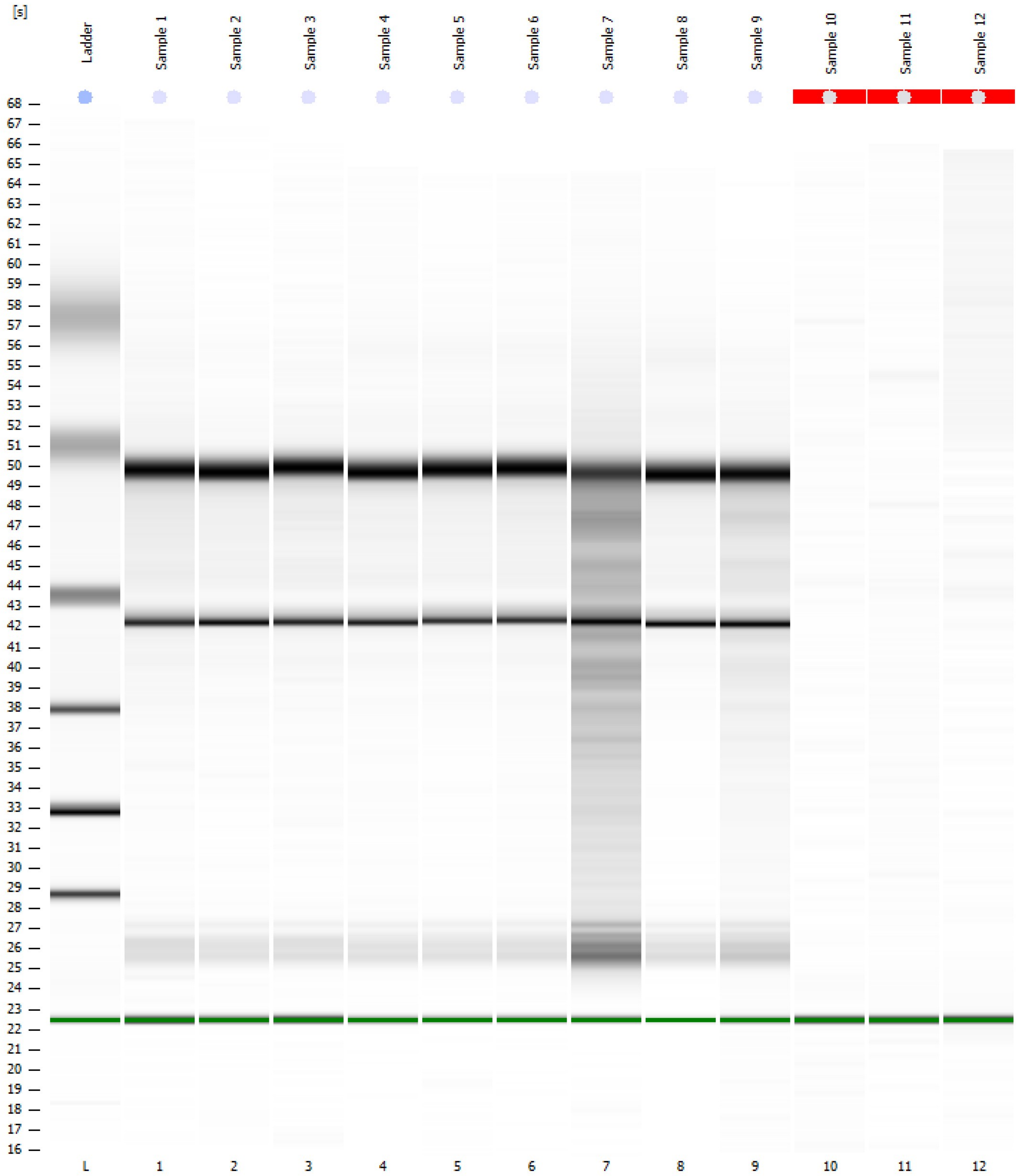


 © Copyright 2003 - 2018 Agilent, Inc. 

### Curves

#### Standard Curve

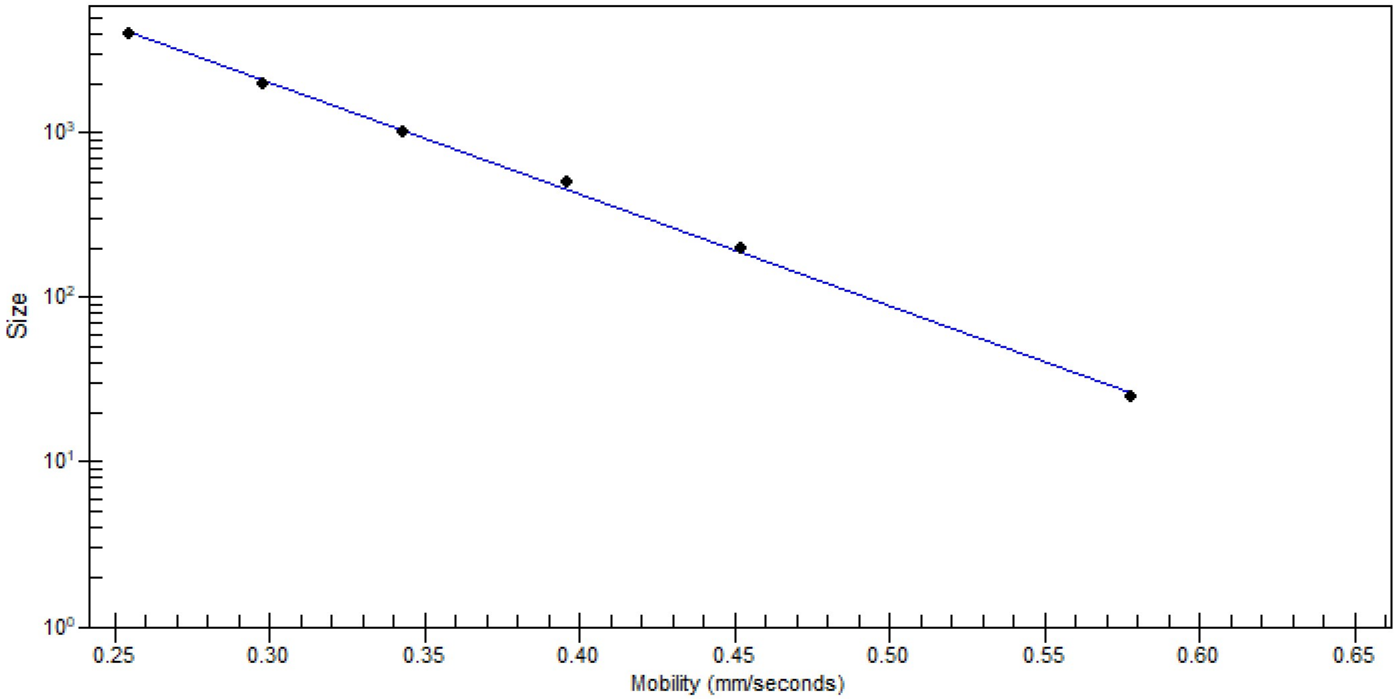


 © Copyright 2003 - 2018 Agilent, Inc. 

Assay Class: Eukaryote Total RNA Nano
Data Path: C:\...Eukaryote Total RNA Nano_DEDAE01485_2022-06-14_15-36-23.xad
Created: 6/14/2022 3:36:23 PM
Modified: 6/14/2022 4:08:38 PM

### Run Logbook

| Description | Number | Source | Category | Sub Category | Time | Time Zone | User | Host |
|---|---|---|---|---|---|---|---|---|
| Run ended on port 1 (Number of wells acquired: 13) | | Instrument | Run | | 6/14/2022 4:00:13 PM | (GMT +02:00) W. Europe Standard Time | user_agilent | FMP1 |
| Run started on port 1 (File: C:\Program Files (x86)\Agilent\2100 bioanalyzer\2100 expert\Data\2022-06-14\2100 expert_Eukaryote Total RNA Nano_DEDAE01485_2022-06-14_15-36-23.xad) | | Instrument | Run | | 6/14/2022 3:36:28 PM | (GMT +02:00) W. Europe Standard Time | user_agilent | FMP1 |
| Product Number : G2939B | | Instrument | Run | | 6/14/2022 3:36:28 PM | (GMT +02:00) W. Europe Standard Time | user_agilent | FMP1 |
| Name : | | Instrument | Run | | 6/14/2022 3:36:28 PM | (GMT +02:00) W. Europe Standard Time | user_agilent | FMP1 |
| Vendor : Agilent Technologies | | Instrument | Run | | 6/14/2022 3:36:28 PM | (GMT +02:00) W. Europe Standard Time | user_agilent | FMP1 |
| Serial# : DEDAE01485 | | Instrument | Run | | 6/14/2022 3:36:28 PM | (GMT +02:00) W. Europe Standard Time | user_agilent | FMP1 |
| Firmware : C.01.069 | | Instrument | Run | | 6/14/2022 3:36:28 PM | (GMT +02:00) W. Europe Standard Time | user_agilent | FMP1 |
| Cartridge : Electrode | | Instrument | Run | | 6/14/2022 3:36:28 PM | (GMT +02:00) W. Europe Standard Time | user_agilent | FMP1 |

 © Copyright 2003 - 2018 Agilent, Inc. 
