## Supplementary material for "*In situ* quantification of ribosome number by electron tomography": Table 1

**Table 1.** Overview of all created electron tomographic data sets of hTERT-RPE-1 cells and *C. elegans* used for detailed analysis.

| **Sample** | **Condition** | **Data set name** |
| --- | --- | --- |
| hTERT-RPE-1 cell, interphase | wild type | [T0611_CRL-4000_WT_Block01_Cell01_TOMO_Interphase01](https://omero.med.tu-dresden.de/webclient/?show=dataset-1715) |
| hTERT-RPE-1 cell, interphase | wild type | [T0661_CRL-4000_WT_Block06_Cell03_TOMO_Interphase02](https://omero.med.tu-dresden.de/webclient/?show=dataset-1718) |
| hTERT-RPE-1 cell, interphase | wild type | [T0661_CRL-4000_WT_Block06_Cell01_TOMO_Interphase03](https://omero.med.tu-dresden.de/webclient/?show=dataset-1713) |
| hTERT-RPE-1 cell, interphase | wild type | [T0661_CRL-4000_WT_Block06_Cell02_TOMO_Interphase04](https://omero.med.tu-dresden.de/webclient/?show=dataset-1714) |
| hTERT-RPE-1 cell, interphase | wild type | [T0661_CRL-4000_WT_Block02_Cell01_TOMO_Interphase05](https://omero.med.tu-dresden.de/webclient/?show=dataset-1717) |
| hTERT-RPE-1 cell, interphase | wild type | [T0661_CRL-4000_WT_Block01_Cell01_TOMO_Interphase06](https://omero.med.tu-dresden.de/webclient/?show=dataset-1716) |
| C. elegans, gonad distal | control(RNAi) | [T0672_N2_control RNAi_Distal Gonad_Worm02_TOMO_Gonad01](https://omero.med.tu-dresden.de/webclient/?show=dataset-1739) |
| C. elegans, gonad distal | control(RNAi) | [T0672_N2_control RNAi_Distal Gonad_Worm08_TOMO_Gonad02](https://omero.med.tu-dresden.de/webclient/?show=dataset-1734) |
| C. elegans, gonad distal | control(RNAi) | [T0672_N2_control RNAi_Distal Gonad_Worm09_TOMO_Gonad03](https://omero.med.tu-dresden.de/webclient/?show=dataset-1727) |
| C. elegans, gonad distal | control(RNAi) | [T0672_N2_control RNAi_Distal Gonad_Worm04_TOMO_Gonad04](https://omero.med.tu-dresden.de/webclient/?show=dataset-1723) |
| C. elegans, gonad pachytene | control(RNAi) | [T0672_N2_control RNAi_Pachytene Gonad_Worm04_TOMO_Gonad01](https://omero.med.tu-dresden.de/webclient/?show=dataset-1732) |
| C. elegans, gonad pachytene | control(RNAi) | [T0672_N2_control RNAi_Pachytene Gonad_Worm07_TOMO_Gonad02](https://omero.med.tu-dresden.de/webclient/?show=dataset-1737) |
| C. elegans, gonad pachytene | control(RNAi) | [T0672_N2_control RNAi_Pachytene Gonad_Worm09_TOMO_Gonad03](https://omero.med.tu-dresden.de/webclient/?show=dataset-1725) |
| C. elegans, gonad pachytene | control(RNAi) | [T0672_N2_control RNAi_Pachytene Gonad_Worm06_TOMO_Gonad04](https://omero.med.tu-dresden.de/webclient/?show=dataset-1740) |
| C. elegans, vulva cell | control(RNAi) | [T0672_N2_control RNAi_Vulva cell_Worm06_TOMO_Interphase01](https://omero.med.tu-dresden.de/webclient/?show=dataset-1733) |
| C. elegans, vulva cell | control(RNAi) | [T0672_N2_control RNAi_Vulva cell_Worm07_TOMO_Interphase02](https://omero.med.tu-dresden.de/webclient/?show=dataset-1735) |
| C. elegans, vulva cell | control(RNAi) | [T0672_N2_control RNAi_Vulva cell_Worm09_TOMO_Interphase03](https://omero.med.tu-dresden.de/webclient/?show=dataset-1729) |
| C. elegans, vulva cell | control(RNAi) | [T0672_N2_control RNAi_Vulva cell_Worm04_TOMO_Interphase04](https://omero.med.tu-dresden.de/webclient/?show=dataset-1722) |
| C. elegans, gonad distal | rpoa-1(RNAi) | [T0675_N2_RPOA-1 RNAi_Distal Gonad_Worm05_TOMO_Gonad01](https://omero.med.tu-dresden.de/webclient/?show=dataset-1738) |
| C. elegans, gonad distal | rpoa-1(RNAi) | [T0671_N2_RPOA-1 RNAi_Distal Gonad_Worm11_TOMO_Gonad02](https://omero.med.tu-dresden.de/webclient/?show=dataset-1730) |
| C. elegans, gonad distal | rpoa-1(RNAi) | [T0671_N2_RPOA-1 RNAi_Distal Gonad_Worm13_TOMO_Gonad03](https://omero.med.tu-dresden.de/webclient/?show=dataset-1724) |
| C. elegans, gonad distal | rpoa-1(RNAi) | [T0679_N2_RPOA-1 RNAi_Distal Gonad_Worm04_TOMO_Gonad04](https://omero.med.tu-dresden.de/webclient/?show=dataset-1743) |
| C. elegans, gonad pachytene | rpoa-1(RNAi) | [T0679_N2_RPOA-1 RNAi_Pachytene Gonad_Worm10_TOMO_Gonad01](https://omero.med.tu-dresden.de/webclient/?show=dataset-1742) |
| C. elegans, gonad pachytene | rpoa-1(RNAi) | [T0671_N2_RPOA-1 RNAi_Pachytene Gonad_Worm11_TOMO_Gonad02](https://omero.med.tu-dresden.de/webclient/?show=dataset-1726) |
| C. elegans, gonad pachytene | rpoa-1(RNAi) | [T0671_N2_RPOA-1 RNAi_Pachytene Gonad_Worm13_TOMO_Gonad03](https://omero.med.tu-dresden.de/webclient/?show=dataset-1741) |
| C. elegans, vulva cell | rpoa-1(RNAi) | [T0675_N2_RPOA-1 RNAi_Vulva cell_Worm05_TOMO_Interphase01](https://omero.med.tu-dresden.de/webclient/?show=dataset-1721) |
| C. elegans, vulva cell | rpoa-1(RNAi) | [T0671_N2_RPOA-1 RNAi_Vulva cell_Worm08_TOMO_Interphase02](https://omero.med.tu-dresden.de/webclient/?show=dataset-1720) |
| C. elegans, vulva cell | rpoa-1(RNAi) | [T0671_N2_RPOA-1 RNAi_Vulva cell_Worm11_TOMO_Interphase03](https://omero.med.tu-dresden.de/webclient/?show=dataset-1736) |
| C. elegans, overview | wild type | [T0596_N2_wild-type_Overview_WormH_TEM_Gonad](https://omero.med.tu-dresden.de/webclient/?show=dataset-1728) |
| C. elegans, overview | rpoa-1(RNAi) | [T0679_N2_RPOA-1 RNAi_Overview_Worm05_TEM_Gonad](https://omero.med.tu-dresden.de/webclient/?show=dataset-1731) |
